# Pan-cancer mapping of single T cell profiles reveals a TCF1:CXCR6-CXCL16 regulatory axis essential for effective anti-tumor immunity

**DOI:** 10.1101/2021.10.31.466532

**Authors:** Livnat Jerby-Arnon, Katherine Tooley, Giulia Escobar, Gitanjali Dandekar, Asaf Madi, Ella Goldschmidt, Conner Lambden, Orit Rozenblatt-Rosen, Ana C. Anderson, Aviv Regev

**Author notes:** Genentech, 1 DNA Way, South San Francisco, CA. These authors contributed equally to this work (Livnat Jerby-Arnon, Katherine Tooley). These authors contributed equally to this work (Ana C. Anderson, Aviv Regev). Correspondence (A.C.A.), (A.R.), (L.J.A.).

## Abstract

Unleashing cytotoxic CD8^+^ T cells for effective cancer treatment requires understanding T cell states across different tumor microenvironments. Here, we developed an algorithm to recover both shared and tumor type specific programs and used it to analyze a scRNA-seq compendium of 38,852 CD8^+^ T cells from 141 patients spanning nine different human cancers. We uncovered a pan-cancer T cell dysfunction program that was predictive of clinical responses to immunotherapy and highlighted CXCR6 as a pan-cancer marker of dysfunctional T cells. In mouse models, CXCR6 increased following checkpoint blockade and was repressed by TCF1. Its ligand, CXCL16, was expressed primarily by myeloid cells, and was co-regulated with antigen presentation genes. CXCR6 deletion decreased Tox, CX3CR1, and Bcl2 expression, predisposing dysfunctional PD-1^+^Tim3^+^CD8^+^ T cells to apoptosis, and compromising tumor growth control. Our approach discovered a TCF1:CXCR6-CXCL16 regulatory axis essential for effective anti-tumor immunity, revealing a new perspective on T cell dysfunction and new opportunities for modulating this cell state.

## INTRODUCTION

CD8^+^ T cells in tumor tissue span a spectrum of functional states, reflecting their stemness, memory potential, effector function, and dysfunction or exhaustion status, whose extent and nature are shaped by the tumor microenvironment (TME). TMEs vary widely within and between cancer types, with variable amounts of vascularization, infiltration of immune cell subsets such as CD4^+^ T helper cells, T regulatory cells, and tumor-associated myeloid cells, and presence of non-immune cell types like cancer-associated fibroblasts (CAFs). These factors influence antigen presentation, immuno-regulatory signals, as well as availability of oxygen and other nutrients, all of which impact CD8^+^ T cell responses.

Harnessing the effector potential of tumor infiltrating CD8^+^ T cells (TILs) has been a key focus of cancer therapeutics. A foremost strategy has been the blockade of immune checkpoint receptors that are expressed at high levels on dysfunctional CD8^+^ TILs. Immune checkpoint blockade (ICB) has been associated with improved CD8^+^ effector function in both pre-clinical cancer models (Woo et al., 2012) and in patients (2–5), where it has achieved durable clinical responses in certain cancers, including melanoma, lung, and renal cancer. Despite the great success of ICB, only 12% of patients across all cancer indications are estimated to respond to ICB (6). The TME has been shown to be a key determinant of ICB response (7–9), but many of the features that determine the relationship between the TME and T cell functionality remain unknown, especially given the variation between tumor types. A greater understanding of CD8^+^ T cell states across multiple tumor types with varied TMEs is critical for informing the development of therapies with the potential to reach a greater number of patients.

Single-cell genomics has allowed us to study T cell states at an unprecedented resolution, yet, to date, most single-cell profiling studies have been limited to small numbers of tumors from a single cancer type (10–19), making it difficult to identify generalizable patterns and avoid overfitting to idiosyncratic or even technical signals. Moreover, because dysfunction and activation are closely intertwined (20–22), it is challenging to decouple them analytically. To pinpoint the most generalizable and novel markers of cell states of interest (*e.g.*, dysfunctional T cells), and functionally investigate the most generalizable “hits”, we need rigorous statistical and machine learning frameworks that can extract and decouple (latent) expression programs across large numbers of cells and patients.

Here, we developed a dimensionality reduction algorithm to identify pan-cancer expression programs within a cell subset and applied it to study CD8^+^ T cells across 141 tumor samples profiled by scRNA-seq from 9 tumor types: melanoma, sarcoma, glioblastoma, breast, lung, liver, pancreatic, ovarian, and colon cancer. We uncovered a pan-cancer T cell dysfunction program decoupled from T cell activation comprised of known and novel T cell dysfunction genes and showed that it is predictive of clinical responses to CTLA-4 and PD-1 blockade in melanoma patients. Among the top 10 hits were the immune checkpoints *CTLA4*, *PD1*, and *TIM3*, as well as *CXCR6*. Investigating the regulatory circuits centered around CXCR6, we found that it was increased upon ICB and directly repressed by TCF1, a transcription factor critical for the maintenance of stem-like CD8^+^ TILs (23, 24). CXCR6 deletion shifted the balance between survival and apoptosis in dysfunctional tumor-antigen specific CD8^+^ TILs, compromising their ability to control tumor growth. We thus uncovered a TCF1:CXCR6-CXCL16 regulatory axis that influences the CD8^+^ T cell differentiation trajectory, ultimately impacting the maintenance of dysfunctional cells and tumor outcome. Our approach opens novel avenues for studying and targeting T cell states and other cell states across a wide range of cancer types.

## RESULTS

### GDMF: Generalizable matrix decomposition framework to recover shared programs across studies

The rapid increase in the number of published single-cell profiling studies opens new opportunities to map cell states across dozens or hundreds of patients. However, in contrast to supervised statistical models that identify differentially expressed genes, which can be designed to account for confounders, covariates, and hierarchical data structures, unsupervised methods for extracting latent patterns representing gene programs often do not model this complexity, making them challenging for application across diverse datasets, as patterns can reflect the confounder rather than shared underlying biology.

To address this challenge, we developed the generalizable matrix decomposition framework (GMDF) for unsupervised meta-analysis of large and diverse datasets. Given a set of scRNA-seq studies, GDMF identifies a low dimensional representation of cell states across conditions by decomposing cell profiles to shared and context-specific metagenes or programs. Going beyond existing methods, GMDF is highly generalizable, such that a user can incorporate nested or hierarchical covariates and contexts in the formulation and implementation of the dimensionality reduction task. “Context” is defined by the user based on the specific data collection and research question, and can denote, for example, a specific cancer type, sex, treatment status, cohort, etc.

GMDF consists of two main steps. The first is formulated as a regularized, non-convex optimization problem that minimizes the reconstruction error (Fig. 1A) and is solved using block coordinate descent. In its second step, GMDF applies a consensus approach, where multiple solutions obtained with different subsamples of the data and initializations are aggregated to a single robust consensus solution (**STAR Methods,** Fig. 1B, Fig. S1A). This second step allows GMDF to mitigate the risk of local minima, non-unique solutions, and overfitting.

**Figure 1.**
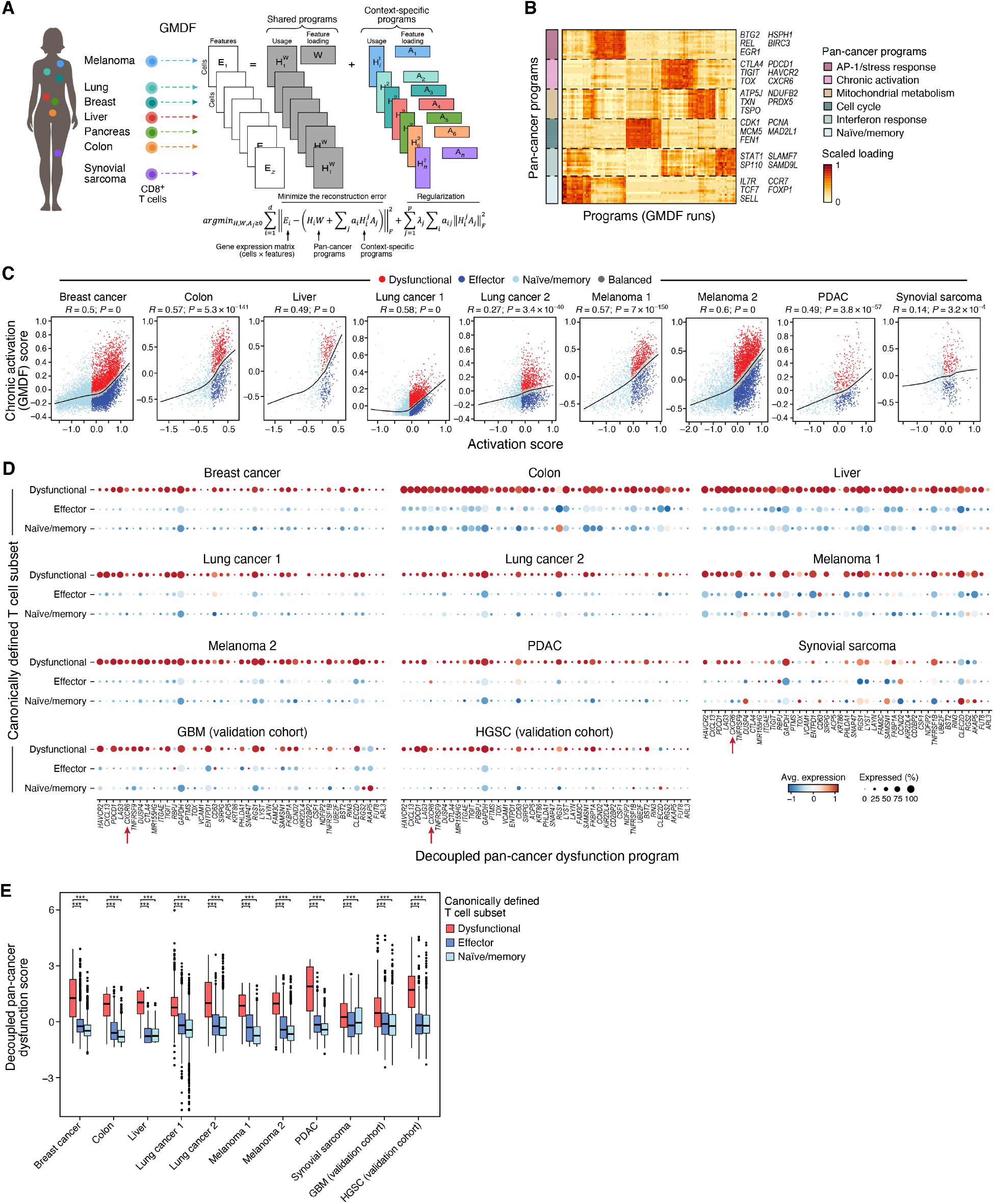
Pan-cancer approach reveals highly generalizable expression programs of T cell dysfunction. **(A)** Analysis approach. scRNA-Seq data of tumor-infiltrating CD8^+^ T cells from seven human cancers (left) were analyzed with GMDF (center) revealing shared (pan-cancer, grey) and context (tumor) specific (color) expression programs (right). **(B)** GMDF identified 6 pan-cancer expression programs in T cells. Top 15 genes (rows) of each program (color-bar on the left) and their weights (W matrix) across programs from the different GMDF solutions (columns). Right: Five representative genes from each program. **(C)** CD8^+^ T cells are stratified by Chronic Activation (GMDF) *vs*. Activation scores. Chronic activation score (y axis, **STAR Methods**) and activation score (x axis, **STAR Methods)** for each CD8 T cell profile (dot) from each of 9 studies (panels) with cells classified as dysfunctional (red), effector (dark blue) or naïve/memory (light blue) by their signatures, or as “balanced” (grey), if their chronic activation score is at the expected level based on the black regression line. Black line: LOWESS regression. Pearson’s R and association P value are marked at the top. **(D,E)** Decoupled pan-cancer dysfunction program distinguishes canonically-defined dysfunctional T cells across tumors. **(D)** Mean expression (dot color, color bar) and proportion of expressing cells (dot size) of genes (columns) in the decoupled pan-cancer T cell dysfunction program in CD8^+^ T cells from 11 tumor scRNA-seq studies stratified by expression of canonical markers (naïve/memory: *CCR7, TCF7, LEF1, SELL;* effector: *NKG7, CCL4, CST7, PRF1, GZMA, GZMB, IFNG, CCL3;* dysfunctional: *PDCD1, TIGIT, HAVCR2, LAG3, CTLA4*). Genes are rank ordered based on their association with the exhausted state (combined p-value). **(E)** Distribution of Overall Expression scores (y axis, **STAR Methods**) of the decoupled pan-cancer dysfunction program (after removing immune checkpoints, i.e., *PDCD1, TIGIT, HAVCR2, LAG3, CTLA4*) in CD8^+^ T cells stratified as in (D). Middle line: median; box edges: 25^th^ and 75^th^ percentiles, whiskers: most extreme points that do not exceed ±IQR*1.5; further outliers are marked individually. ***p < 0.001, mixed effects test.

In contrast to other methods, such as integrative-NMF (iNMF) (25) and LIGER (26), GMDF does not force any coupling of the shared and context-specific programs (**STAR Methods**), and thus outperforms previous methods in this task, as we show using simulated data (Fig. S1B-E). Specifically, while LIGER and iNMF identify shared programs with context-specific extensions, GMDF provides sufficient flexibility in identifying both latent shared as well as context-specific programs without enforcing any type of coupling between them. Comparing these methods (Fig. S1B-E**, STAR Methods**), we find that enforcing a specific structure, as done by LIGER, results in suboptimal decompositions and in misclassification of context-specific programs (*i.e.*, that occur only in a specific dataset) as shared programs.

### GDMF identifies pan-cancer T cell expression programs across tumor types

To map CD8^+^ T cell states across cancer types, tissues, and microenvironments, we assembled a pan-cancer scRNA-seq compendium that integrates nine scRNA-Seq datasets, consisting of 33,161 intra-tumoral CD8^+^ T cells, collected from 134 cancer patients with melanoma, sarcoma, pancreatic, liver, colon, lung, and breast cancer (10–14,16–19). We applied GMDF to this pan-cancer CD8 T cell compendium with cohort covariates (**STAR Methods**), to identify shared (“pan-cancer”) programs.

GMDF identified six pan-cancer programs that were shared across all nine cohorts of the pan-cancer CD8 T cell compendium, which we annotated as AP-1/stress response, mitochondrial metabolism, cell cycle, interferon response, naive/memory, and chronic activation (Fig. 1B, **Tables S1 and S2**). The chronic activation program contained genes associated with T cell dysfunction, including all known immune checkpoints (*TIM3*, *TIGIT*, *PD1*, *CTLA4*, and *LAG3*) and the transcription factor *TOX* (27–29). In line with previous studies (11, 22), it also included effector genes (*GZMB, IFNG, FASLG, ZBED2*), reflecting the coupling between T cell dysfunction and activation (20–22).

To further examine the dysfunction-activation association, we computed an “Activation score” for each cell based on its Overall Expression (11) (**STAR Methods**) of canonical effector function markers (*i.e.*, *NKG7, CCL4, CST7, PRF1, GZMA, GZMB, IFNG, CCL3*) minus its Overall Expression of canonical naïve/memory T cell markers (*i.e.*, *CCR7, TCF7, LEF1, SELL*). Naïve cells, with high expression of the naïve markers and low expression of the effector markers, were all characterized by low expression of the pan-cancer chronic activation program identified here. However, as cells became more activated, they were associated with increasingly higher values of the chronic activation program and showed more inter-cellular variation (Fig. 1C). To further decouple activation and dysfunction scores, we normalized the chronic activation score based on the expected value given the “Activation score” (LOWESS regression line shown in black, Fig. 1C, **STAR Methods**). Next, we annotated each T cell as naïve/memory (low activation scores), effector (high activation and lower-than-expected chronic activation scores, *i.e.*, below the LOWESS regression line), or dysfunctional (high activation and higher-than-expected chronic activation scores, *i.e.*, above the LOWESS regression line, (**STAR Methods,** Fig. 1C). Finally, using these three cell annotations and a multilevel regression modeling framework (**STAR Methods**), we identified the unique features of each of these cell states across all cohorts, resulting in three “decoupled pan-cancer programs” of Effector, Dysfunction, and Naïve/Memory cell states (Fig. 1D and **S2A,B**, **Table S3**). The resulting decoupled pan-cancer dysfunction program consisted of 43 genes (Fig. 1D), including the established checkpoints *PD1*, *CTLA4*, and *TIM3* among the top 10.

To test the generalizability of these programs we used the canonical effector and naïve/memory markers (listed above), together with immune checkpoint genes (*PDCD1, TIGIT, HAVCR2, LAG3, CTLA4*) to first stratify the cells, and then examined whether using only the *de novo* genes in the three decoupled pan-cancer programs (*i.e.*, after removing any of the canonical markers) could identify the pertaining cell population. Indeed, based on this analysis, each of these *de novo* programs marked the respective T cell populations across all cohorts, as well as in two independent scRNA-seq studies (not included in the compendium) of glioblastoma (15) and of ovarian cancer (30) (P < 8.32*10^-22^, 1.2*10^-16^, 1.33*10^-46^, mixed-effects test, for the dysfunction, effector, and naïve/memory programs in glioblastoma, respectively, and similarly, P < 4.95*10^-36^, 4.57*10^-3^, 1.1*10^-10^, in ovarian cancer, **Methods**, Fig. 1E **and** Fig. S2C-D).

### The decoupled pan-cancer T cell dysfunction program predicts clinical response to immunotherapy

To test the clinical relevance of the three decoupled pan-cancer programs as well as the pan-cancer cell cycle program, we assessed their expression in scRNA-seq data of CD8^+^ TILs collected from melanoma patients before and after ICB treatment (17). Each of the programs highlighted distinct populations of cells (Fig. 2A). The decoupled pan-cancer dysfunction program scored very highly in cells from non-responder patients (Fig. 2A, P = 5.70*10^-30^, mixed-effects test; pre-and post-treatment combined). This was in accordance with the previously reported (17) correlation of *ENTPD1* and *HAVCR2*, both contained in the decoupled pan-cancer dysfunction program, with failure to respond to ICB (Fig. S3A,B). Interestingly, the decoupled pan-cancer dysfunction program also includes *SIRPG* (Fig. S3C), the ligand of *CD47*, consistent with previous studies linking CD47 expression on cancer cells to T cell function (31–33) and lack of ICB response (11). In addition, the cell cycle program scored positively in cells from non-responder patients both pre-and post-treatment (P = 4.23*10^-9^, respectively, mixed-effects), while the decoupled naïve/memory program, containing *TCF7* (Fig. S3D), showed the opposite trend (P = 6.01*10^-21^, mixed-effects), both before and after treatment.

**Figure 2.**
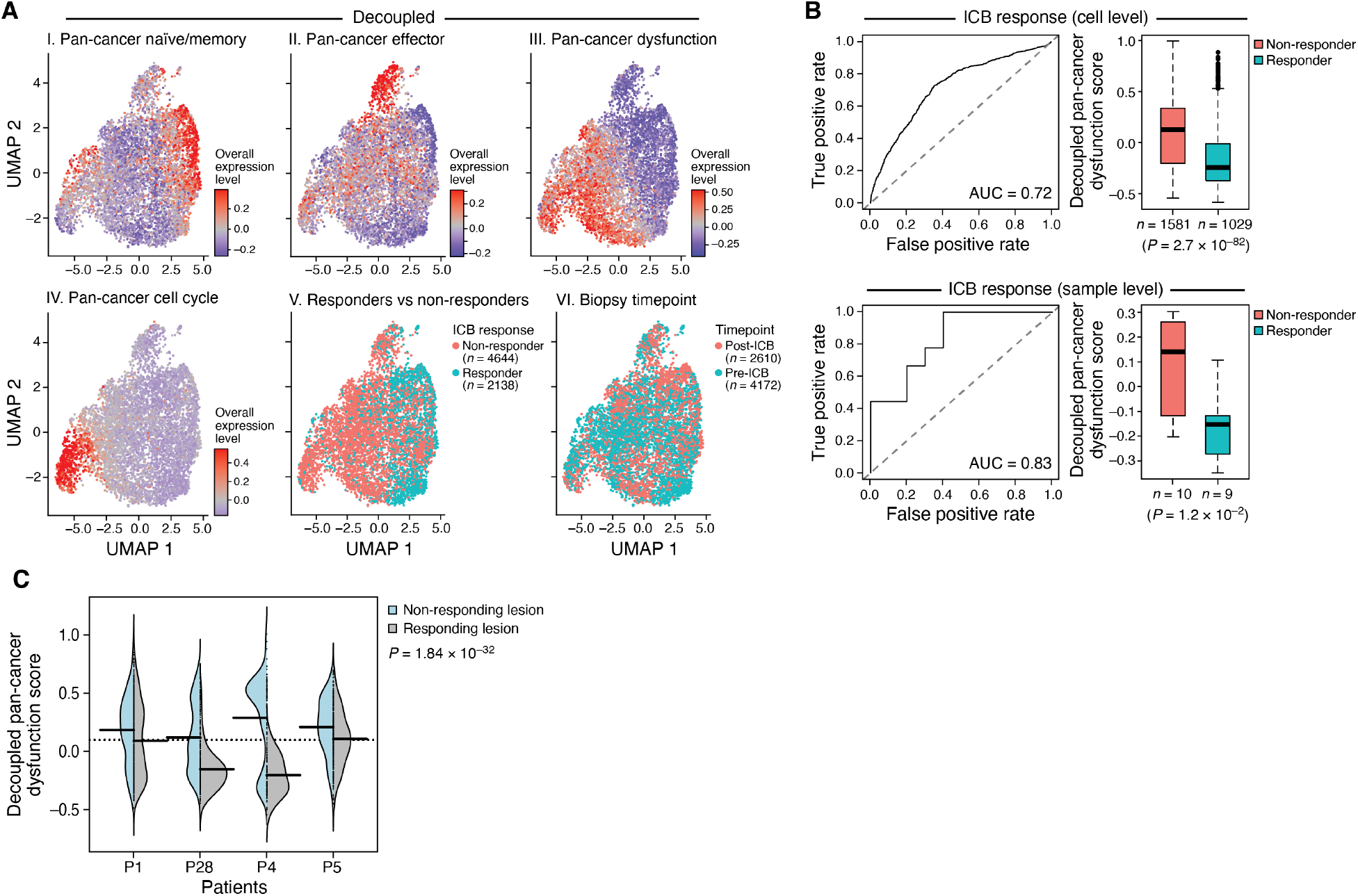
The pan-cancer T cell dysfunction program predicts ICB responses. **(A)** Pan cancer expression program is associated with CD8 T cells from non-responders. Uniform Manifold Approximation and Projection (UMAP) embedding of CD8 T cell profiles (dots) from 48 melanoma tumors (17), colored by the Overall Expression of the decoupled pan-cancer naïve/memory (I), effector (II), and dysfunction (III), or pan-cancer cell cycle (IV) program, ICB response status (V), or pre-versus post-treatment biopsy status (VI). **(B)** Pan-cancer T cell dysfunction program is predictive of response to ICB. Left: True positive (y axis) and false positive (x axis) rates when predicting the clinical response of the tumor based on different levels of the pan-cancer T cell dysfunction program score at either the CD8^+^ T cell (top) or sample (bottom) level. Right: Distribution of the Overall Expression (y axis) of the decoupled pan-cancer dysfunction program in responders (red) or non-responders (blue) at the cell (top) and sample (bottom) level. Middle line: median; box edges: 25^th^ and 75^th^ percentiles, whiskers: most extreme points that do not exceed ±IQR*1.5; further outliers are marked individually. **(C)** Higher level of the pan-cancer T cell dysfunction program in non-responding lesions. Distribution of Overall Expression of the decoupled pan-cancer dysfunction program (**STAR Methods**) in the responding (grey) and non-responding (light blue) lesions of patients (x axis) with mixed responses.

The expression of the decoupled pan-cancer dysfunction program in CD8^+^ TILs pre-treatment was predictive of clinical ICB responses (AUC = 0.72 and 0.83, when examined at the single-cell, P = 2.7*10^-82^, and sample-level, P = 0.012, t-test, Fig. 2B), and was not induced in the post-ICB samples (P>0.1). Interestingly, some of the patients in the cohort had mixed responses, with both responding and non-responding lesions. The decoupled pan-cancer dysfunction program predicted response at the lesion level in these patients, such that it was significantly more highly expressed in cells from the non-responding *vs*. responding lesions in four different patients (P < 1*10^-30^, Fisher combined test, Fig. 2C; the responding and non-responding lesions obtained from the same patient were not always sampled at the same time).

### CXCR6 is a high-ranking gene in the decoupled pan-cancer dysfunction program

Examining the decoupled pan-cancer dysfunction program genes for novel potential regulators of the dysfunctional T cell state, we observed *CXCR6* among the top 10 program genes, as it had a high loading in the GMDF consensus solution and consistently marked the dysfunctional T cells across all the different cancers examined (Fig. 1D). *CXCR6* has been previously studied in the context of immunization, infection, and inflammation in mice, where it was associated with improved immune responses along with effector and memory T cell recruitment and trafficking (34, 35). Interestingly, in a chronic viral infection model, CXCR6 marked cells labeled as “advanced exhausted” (36).

CXCR6 was predominantly expressed in T cells, and its ligand, CXCL16, was predominantly expressed by myeloid cells, based on our analysis of four tumor types in our pan-cancer scRNA-seq compendium where non-T cells were profiled as well (sarcoma, lung, melanoma, and the glioblastoma validation dataset) (11–13,15,17) (Fig. 3A). Based on these datasets we identified a CXCL16 program (Fig. S3E), consisting of the genes that are co-expressed with *CXCL16* in macrophages. The program included complement genes (*C1QA*, *C1QB, C1QC*), MHC class II genes (*HLA-DMB*, *HLA-DOA*), and the co-stimulatory receptor *CD86*, indicating that CXCL16 marks activated cells engaged in antigen cross-presentation (**Table S4**). Indeed, we found CXCL16 to be co-expressed with antigen processing and presentation signatures in macrophages in all datasets (Fig. 3B). We therefore hypothesized that the CXCR6-CXCL16 axis may positively regulate the dysfunctional T cell state, mediating spatiotemporal control of chronically activated T cells in the TME.

**Figure 3.**
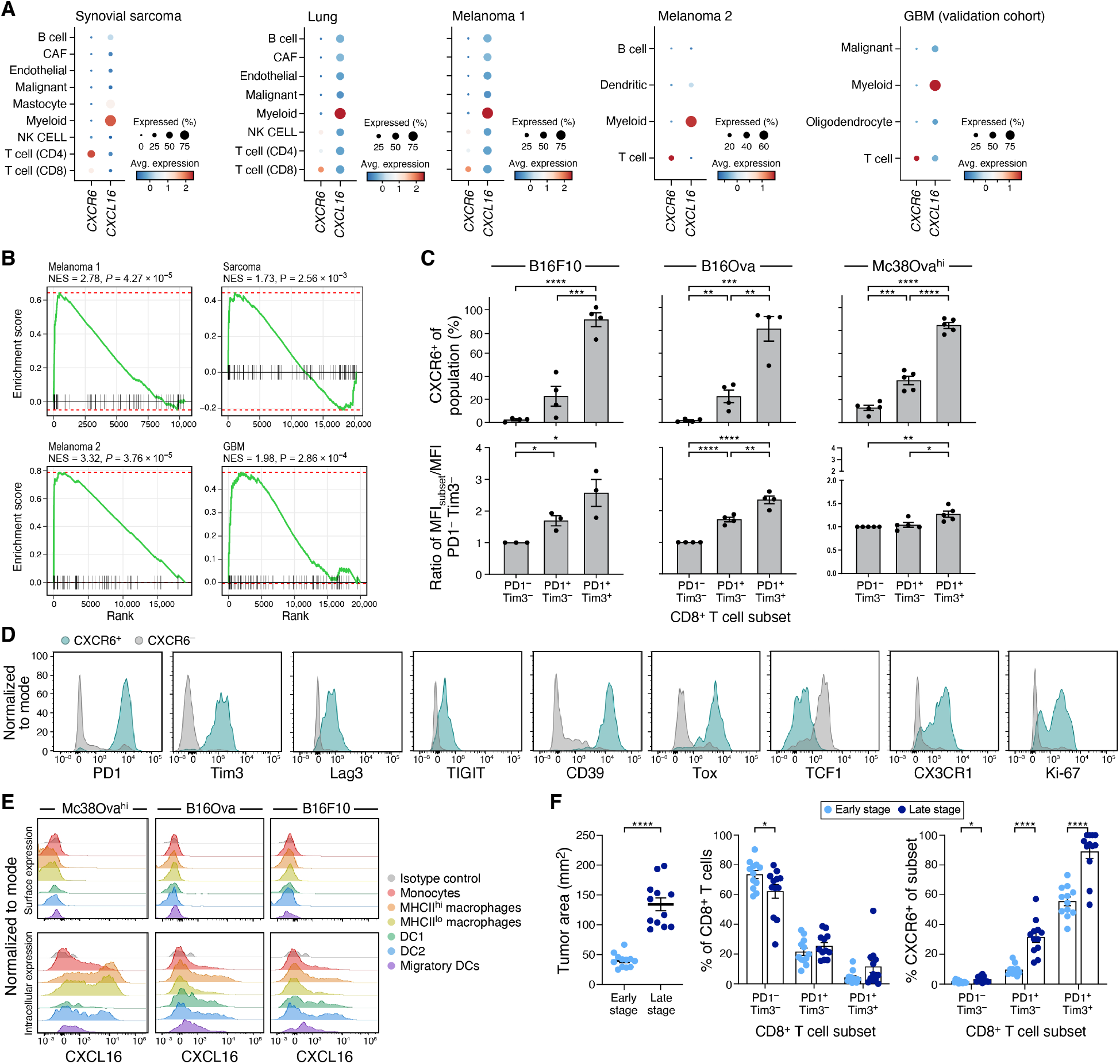
CXCR6 and CXCL16 expression in human and murine tumors track with dysfunctional T cells and myeloid cells respectively. **(A)** CXCR6 and CXCL16 expression in human tumors. Mean expression (dot color, colorbar) and fraction of expressing cells (dot size) for CXCR6 and CXCL16 (columns) across cell types (rows) in different human tumor studies (panels). **(B)** CXCL16 is co-expressed with antigen presentation genes in macrophages. GSEA plots obtained for the KEGG_ANTIGEN_PROCESSING_AND_PRESENTATION gene set when compared against the ranking of genes based on their co-expression with CXCL16 in macrophages in the indicated tumor type. **(C,D,F)** CXCR6 expression tracks with CD8^+^ T cell dysfunction and tumor size in mouse tumors. **(C)** Frequency (y axis, top) and geometric mean fluorescence intensity (MFI, y axis, bottom) of CXCR6 expression in the indicated CD8^+^ TILs populations (x axis) harvested from mice bearing B16F10, B16Ova, or Mc38Ova^Hi^ tumors. **(D)** Representative distributions of expression levels (x axis = fluorescence intensity) on CXCR6^+^ (green) and CXCR6^-^ (grey) CD8^+^ TILs isolated from B16Ova tumors. **(E)** CXCL16 is expressed in myeloid cells in mouse tumors. Representative distributions of CXCL16 surface (top) or intracellular (bottom) expression (x axis) in myeloid cells from Mc38Ova^Hi^, B16Ova, or B16F10 tumors. **(F)** Left: B16Ova tumor size (y axis), mm^2^, at early (light blue) or late (dark blue) tumors. Middle: Percent of CD8^+^ T cells (y axis) belonging to each population defined by PD1-and Tim3 expression (x axis) in early (light blue) or late (dark blue) tumors. Right: Percent of CXCR6 expressing cells belonging to each population defined by PD1-and Tim3 expression (x axis) in early (light blue) or late (dark blue) tumors. Mean ± SEM is shown. *p < 0.05, **p < 0.01, ***p < 0.001, ****p<0.0001, t test.

### CXCR6 expression tracks with T cell dysfunction and tumor progression across different tumor microenvironments in mouse models

To further relate the CXCR6-CXCL16 axis to T cell dysfunction, we assessed CXCR6 expression on CD8^+^ T cells in different murine tumor models that are highly (Mc38 colon carcinoma) or poorly (B16F10 melanoma) immunogenic and vary in their expression of Ova as an ectopic tumor antigen. Across all models, CXCR6 expression increased gradually from PD1^-^Tim3^-^ (comprised of naïve-like cells) to PD1^+^Tim3^-^ (comprised of stem-like and effector-like cells) to PD1^+^Tim3^+^ (terminally dysfunctional) CD8^+^ TILs (Fig. 3C). Compared to CXCR6^-^ counterparts, CXCR6^+^ TILs had higher expression of the proliferative marker Ki-67, CX3CR1 (associated with effector function), along with multiple co-inhibitory receptors (PD1, Tim3, Lag3, Tigit, CD39) and the transcription factor Tox (27–29), all of which are associated with dysfunction (Fig. 3D). Notably, CXCR6^+^ T cells had lower TCF1, indicating concomitant loss of stemness with acquisition of CXCR6 expression (Fig. 3D). In line with the human pan-cancer compendium, CXCL16 was expressed by myeloid cells across the three tumor models, with the highest expression in MHCII^hi^ macrophages, followed by MHCII^lo^ macrophages, DC1, and DC2 subsets (Fig. 3E). This is consistent with a model where the CXCR6-CXCL16 axis could mediate interactions between effector and dysfunctional PD1^+^Tim3^+^CD8^+^ TILs with antigen-presenting cells in the TME.

We further determined how CXCR6 expression changed over the course of tumor progression, comparing CD8^+^ TILs from B16Ova tumors harvested at the early versus late stages of tumor progression. While the proportions of PD1-and Tim3-expressing CD8^+^ T cell subsets remained mostly unchanged, the proportion of CXCR6 expressing cells among both PD1^+^Tim3^-^ and PD1^+^Tim3^+^ cells increased with tumor progression (Fig. 3F). Thus, CD8^+^ T cells increasingly acquired CXCR6 expression as they transitioned to effector and finally to the terminally dysfunctional T cell states with tumor progression.

### CXCR6 is up-regulated following ICB treatment and directly repressed by TCF1

The gradient of increasing expression of CXCR6 from effector to dysfunctional CD8^+^ TILs (Fig. 3C) prompted us to examine whether its expression changed in response to immune checkpoint blockade. In CD8^+^ TILs from melanoma patients collected pre-and post-treatment (17), CXCR6 expression was increased following ICB treatment (P = 4.63*10^-4^, mixed-effects). To test this in a controlled setting, we examined two different tumor models (Mc38Ova^Hi^ and B16Ova) and two different ICB therapies (anti-PD1 and anti-PD-L1+anti-Tim-3).

In the immunogenic Mc38Ova^Hi^ tumor model, the fraction of cells expressing CXCR6 increased with ICB in both the PD1^+^Tim3^-^ and PD1^+^Tim3^+^ CD8^+^ TIL populations (Fig. 4A), whereas in the less immunogenic B16Ova model, the proportion of CXCR6^+^ cells increased upon ICB in the PD1^+^Tim3^-^, but not the PD1^+^Tim3^+^ CD8^+^ TIL populations, where nearly all cells already expressed it prior to ICB (Fig. 4B). The increase in CXCR6 expressing cells upon ICB was most robust across tumor models in the PD-1^+^Tim3^-^ CD8^+^ T cell population that contains both stem-like and effector-like CD8^+^ T cells. In contrast, TCF1, which is required for the maintenance of stem-like CD8^+^ T cells in both cancer and chronic virus infection (23,24,37), is known to decrease in this population upon ICB.

**Figure 4.**
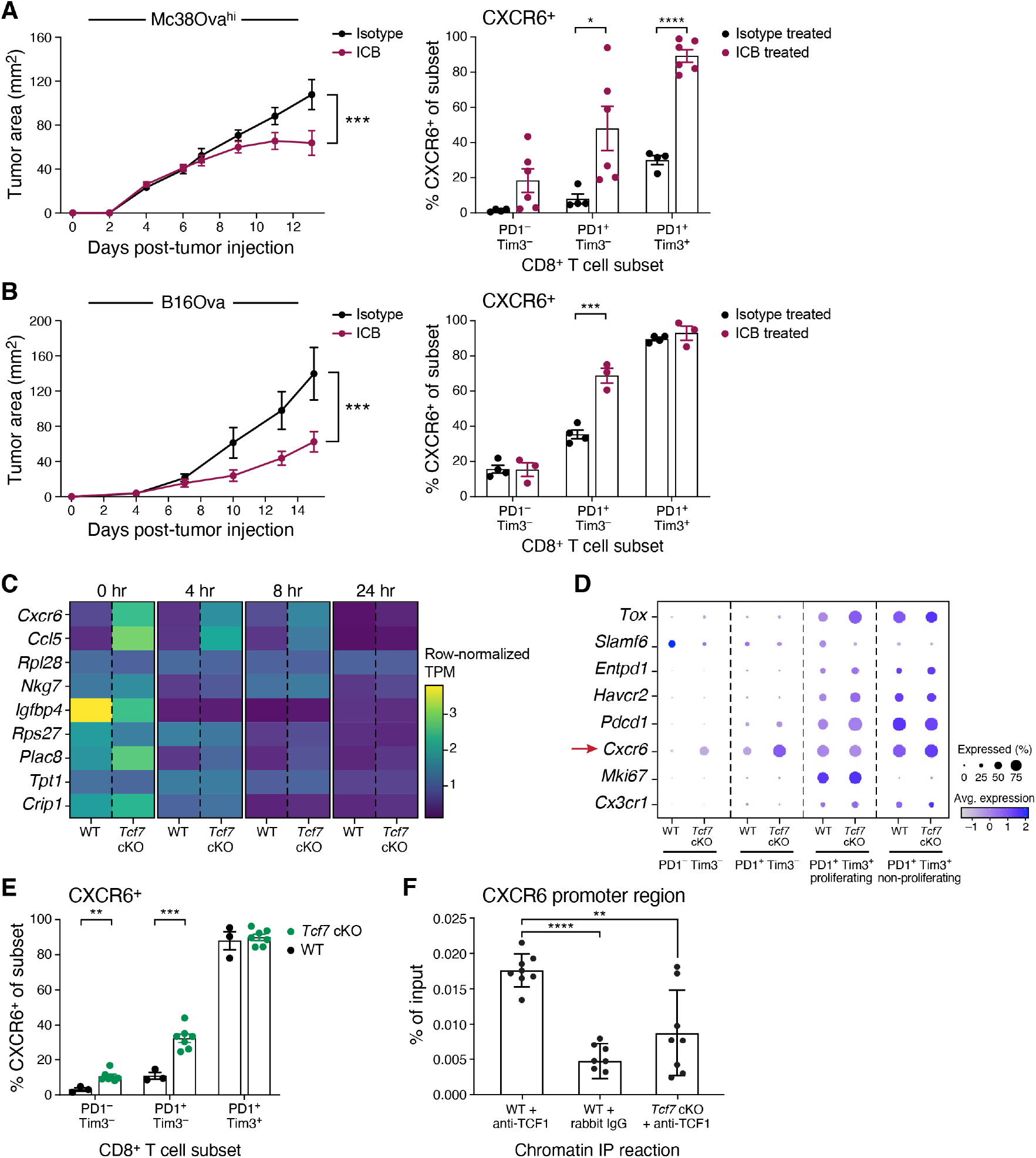
CXCR6 expression is upregulated upon immune checkpoint blockade and repressed by TCF1. **(A,B)** Increased CXCR6 expression in ICB treated mouse tumors. Mean tumor growth (y axis, left, ± SEM) and CXCR6 expression (y axis, right) across indicated CD8^+^ TILs populations (x axis, right) in Mc38Ova^Hi^ **(A)** or B16Ova **(B)** tumors implanted in wild type mice and treated with Anti-PD1 (A, red), Anti-PD-L1 + anti-Tim3 (B, red) or isotype control (black). *p < 0.05, ***p < 0.001. Left: linear mixed model. Right: unpaired t test. **(C-F)** TCF1 represses CXCR6 expression. **(C)** Expression (row-normalized TPM, colorbar) of top differentially expressed genes (rows) between wild type (WT) (E8i-Cre^-^, TCF7^FL/FL^) or *Tcf7* cKO (E8i-Cre^+^, TCF7^FL/FL^) cells at different timepoints after T cell activation (columns). **(D)** Mean expression (dot color, colorbar) and fraction of expressing cells (dot size) of key genes (rows) across T cells from different CD8^+^ T cell clusters (columns, **STAR Methods**) as determined by scRNA-Seq of CD8^+^ T cells from B16Ova tumors implanted in WT or *Tcf7* cKO mice. **(E)** Percent of CXCR6^+^ cells (y axis) in each indicated CD8^+^ TILs population (x axis) from WT (black) or *Tcf7* cKO (green) mice implanted with B16Ova. **p < 0.01, ***p < 0.001, unpaired t test. **(F)** Percent input (y axis) following Chromatin-Immunoprecipitation (ChIP) determined by quantitative PCR for the CXCR6 promoter region on CD8^+^ T cells isolated from wildtype or *Tcf7* cKO mice with anti-TCF1 or rabbit IgG control antibodies (x axis). *p < 0.05, **p < 0.01, ***p < 0.001, ****p<0.0001, paired t test (comparing WT samples) and unpaired t test (comparing WT to *TCF7* cKO).

We therefore hypothesized that TCF1 may have a regulatory relationship with CXCR6. To test this hypothesis, we measured bulk RNA-Seq profiles of CD8^+^ T cells isolated from either wild type (WT) mice or mice bearing conditional deletion of *Tcf7* (the gene encoding TCF1) in mature CD8^+^ T cells (*Tcf7* cKO) over a time course of *in vitro* activation. CXCR6 was among the top 5 differentially expressed genes between WT and *Tcf7* cKO both prior to activation (timepoint 0, P = 6.23*10^-23^, Wald test) and at subsequent timepoints, with the expression difference gradually decreasing over the course of activation (Fig. 4C). Moreover, scRNA-seq profiling of CD8^+^ TILs from WT or *Tcf7* cKO mice bearing B16Ova tumors showed a significant increase in CXCR6 expression in the naïve-like PD1^-^Tim3^-^ population (P < 1*10^-4^, Wilcoxon) and in the PD1^+^Tim3^-^ population (P < 1*10^-4^, Wilcoxon) in *Tcf7* cKO CD8^+^ TILs compared to WT (Fig. 4D), which we confirmed at the protein level (Fig. 4E).

These results suggested that *Tcf7*/TCF1 acted genetically as a repressor of CXCR6 expression, consistent with studies that show that TCF1 can function as a transcriptional repressor (38). Supporting direct regulation, a TCF1 binding event in the CXCR6 locus has been reported in ChIP-seq data (39) in a chromatin region also shown to be open in naïve T cells by ATAC-seq (40) (Fig. S4A). We confirmed TCF1 binding in the CXCR6 locus by ChIP-PCR performed on CD8^+^ T cells from WT and *Tcf7* cKO mice using primers flanking this region (Fig. 4F). Thus, CXCR6 expression tracked with increased effector differentiation and tumor growth control upon ICB and was repressed by *Tcf7/*TCF1, which is known to restrain effector differentiation (41, 42).

### CXCR6 is required to maintain tumor control

To study the role of CXCR6 in determining the function of anti-tumor CD8^+^ T cells, we next utilized CRISPR-Cas9 to delete CXCR6 in mature OTI CD8^+^ T cells, followed by transfer into mice bearing Ova-expressing tumors (Fig. 5A). This approach deletes genes at the time of T cell activation, thus allowing for early T cell activation to proceed normally and bypassing any compensatory mechanisms that can arise when genes are deleted during development. We confirmed that CRISPR CXCR6-KO OTI cells were efficiently deleted *in vivo* (Fig. S4B) and that CRISPR-Control OTI cells elicited tumor growth control in both the B16Ova and Mc38Ova^Hi^ models.

**Figure 5.**
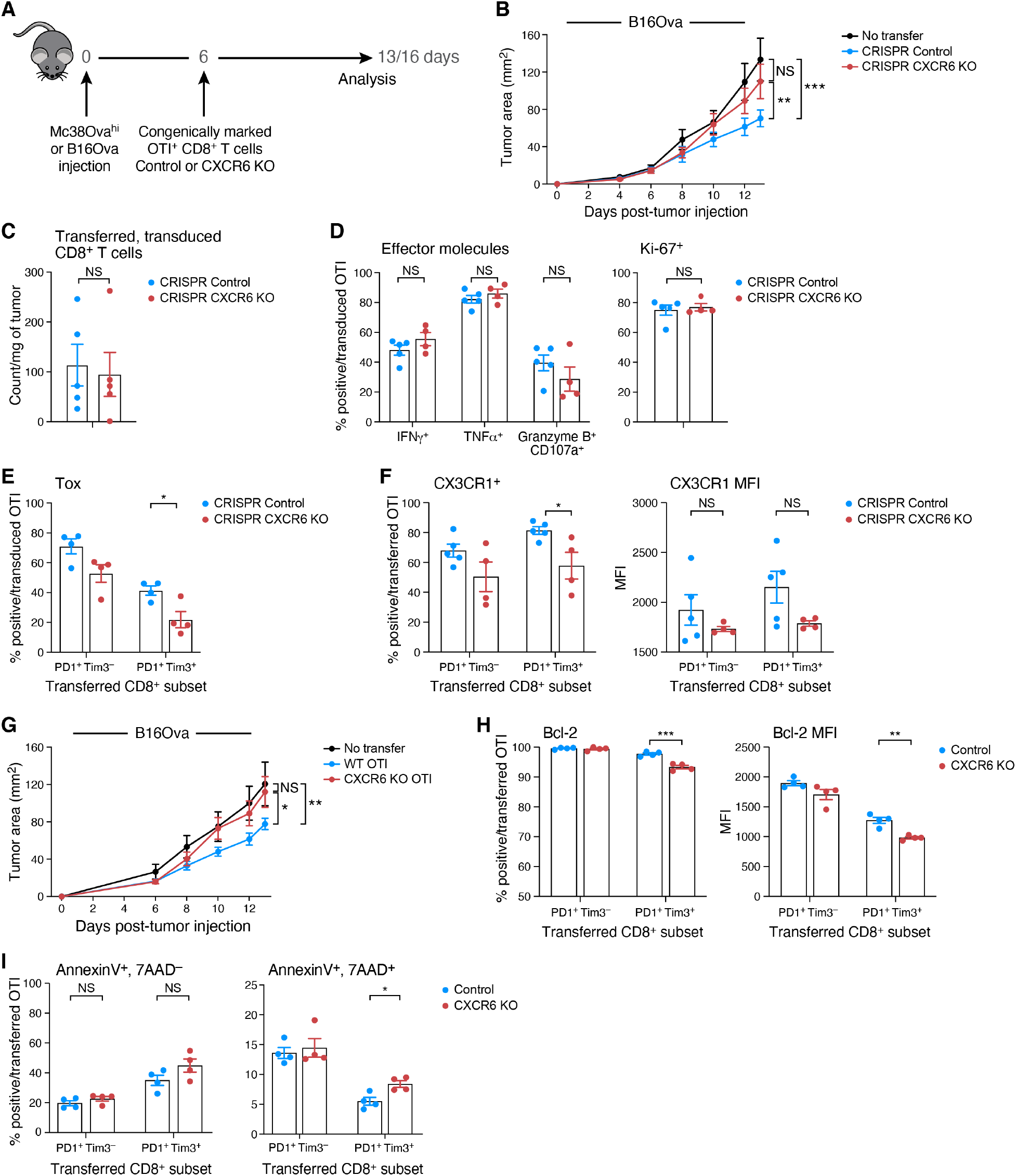
CXCR6 knockout reduces anti-tumor immunity. **(A)** Experimental design for adoptive transfer experiments. **(B)** Reduced tumor control in CXCR6 CRISPR-KO. Tumor size (y axis, mm2, mean ± SEM) over time (x axis, days) for B16Ova in Cas9^+^, CD45.1/.2 mice adoptively transferred with CRISPR CXCR6-KO (red), CRISPR-Control (blue) OT-I T cells, or no transfer control (black). **p < 0.01, ***p < 0.001, linear mixed model. **(C,D)** CXCR6 CRISPR-KO does not impact number and effector features of transferred cells. Number of transferred, transduced cells per mg of tumor tissue (C, y axis, mean ± SEM), frequency of transferred, transduced cells (D, left, y axis, Mean ± SEM) that express IFN*γ*^+^, TNF*α*^+^, or Granzyme B^+^, CD107a^+^ cells (D, left, x axis) after ex-vivo activation with Ova 257-264 peptide, and frequency of Ki-67^+^ cells (D, right, y axis, Mean ± SEM) among CRISPR CXCR6-KO (red), CRISPR-Control (blue) OT-I T cells. **(E)** Reduced Tox expression by CXCR6 CRISPR-KO in PD1^+^Tim3^+^ OTI cells. Frequency of Tox^+^ cells (y axis, mean ± SEM) in transferred, transduced cells within PD1^-^Tim3^+^ and PD1^+^Tim3^+^ CD8^+^ TILs (x axis). *p < 0.05, t test. **(F)** CXCR6 CRISPR-KO, dysfunctional CD8^+^ T cells express lower levels of CX3CR1. Frequency of CX3CR1^+^ cells (left, y axis, Mean ± SEM) and geometric mean fluorescence (MFI) of CX3CR1^+^ cells (right y axis, Mean ± SEM) of transferred, transduced cells within PD1^-^Tim3^+^. *p < 0.05, t test. **(G)** CXCR6 KO cells fail to control tumor growth. Tumor size (y axis, mm2, mean ± SEM) over time (x axis, days) for B16Ova in CD45.1 mice adoptively transferred on day 6 with either CXCR6 knockout (red), control OT-I T (blue) cells, or no transfer control (black). *p < 0.05, **p < 0.01, linear mixed model. **(H,I)** CXCR6 deletion predisposes dysfunctional PD-1^+^Tim3^+^CD8^+^ T cells to apoptosis. **(H)** Frequency of Bcl-2^+^ cells (left, y axis, Mean ± SEM) and level of Bcl-2 in Bcl-2^+^ cells (right, geometric mean fluorescence (MFI), y axis) within PD1^-^Tim3^+^ and PD1^+^Tim3^+^ transferred CD8^+^ TILs (x axis) in B16Ova in CD45.1 mice adoptively transferred on day 6 with either CXCR6 knockout (red), control OT-I T (blue) cells. **(I)** Frequency of Annexin V^+^, 7AAD^-^ (left, y axis, Mean ± SEM) and of Annexin V^+^, 7AAD^+^ (right, Mean ± SEM) transferred cells within PD1^+^Tim3^-^ and PD1^+^Tim3^+^ CD8^+^ TILs (x axis) in B16Ova in CD45.1 mice transferred with either CXCR6 knockout (red), control OT-I T (blue) cells. *p < 0.05, **p < 0.01, ***p < 0.001, t test.

Conversely, CRISPR CXCR6-KO OTI T cells failed to control tumor growth (Fig. 5B and Fig. S4C), such that the tumor growth curve in mice receiving CRISPR CXCR6-KO OTI was indistinguishable from that of mice that received no T cell transfer. We did not observe significant differences in the numbers of transduced cells or in the PD1-and Tim3-expression profile between CRISPR CXCR6-KO OTI or CRISPR-Control OTI (Fig. 5C **and** Fig. S4D,E). Surprisingly, functional analyses showed no major differences in the proliferation, cytokine production, or cytotoxic capacity of CRISPR-Control and CRISPR CXCR6-KO OTI (Fig. 5D). However, Tox expression was reduced, especially on CRISPR CXCR6-KO PD1^+^Tim3^+^ OTI compared to CRISPR-Control PD1^+^Tim3^+^ OTI (Fig. 5E); Tox expression has been shown to impact the survival of CD8^+^ TILs (28). There were also fewer CRISPR CXCR6 KO PD1^+^Tim3^+^ OTI expressing CX3CR1 and a trend towards lower CX3CR1 expression levels compared to CRISPR-Control OTI (Fig. 5F).

CXCR6 also impacted tumor growth in another adoptive transfer model using CD8^+^ T cells from OTI mice crossed with mice in which GFP cDNA is knocked into the endogenous CXCR6 locus disrupting its expression (CXCR6 KO). In line with our results with OTI cells harboring CRISPR-mediated CXCR6 deletion, CXCR6 KO cells completely failed to control tumor growth compared to control cells (Fig. 5G). There was also no significant difference in the number of transferred cells present in the tumor at time of analysis (Fig. S5A) or in the proliferation, cytokine production, and cytotoxicity of transferred cells (Fig. S5B). However, in line with our data in CRISPR CXCR6-KO CD8^+^ T cells, the expression level of CX3CR1 was decreased in CXCR6 KO PD1^+^Tim3^+^ cells compared to control (Fig. S5C, right), although the proportion of CX3CR1^+^ cells was unchanged (Fig. S5C, left). As CX3CR1 expression has been associated with the pro-survival protein Bcl2 (43, 44), we examined if there were differences in Bcl2 between CXCR6 KO and WT cells. While the level of Bcl2 in transferred cells was overall high, the proportion of Bcl2-expressing cells and Bcl2 expression level were significantly lower in PD1^+^Tim3^+^ CXCR6 KO cells compared to control cells (Fig. 5H). Moreover, there were more apoptotic (Annexin V^+^, 7AAD^+^) PD1^+^Tim3^+^ CXCR6 KO cells compared to control, but no major changes in the proportion of pre-apoptotic (Annexin V^+^, 7AAD^-^) cells between CXCR6 KO and WT (Fig. 5I). Thus, these data support a model where CXCR6-CXCL16 interactions regulate the survival of PD1^+^Tim3^+^ cells via alterations in CX3CR1 and Bcl2, such that in the absence of CXCR6, cells are more apoptotic and therefore less effective at mediating tumor clearance.

## DISCUSSION

Here, we devised and applied an integrative approach to uncover shared regulatory mechanisms governing CD8^+^ T cell states across a broad spectrum of human cancers. We identified a novel and generalizable pan-cancer T cell dysfunction program decoupled from T cell activation. One of the top-ranking genes in the program was *CXCR6*, which was negatively regulated by TCF1 and was predominantly expressed in PD1^+^Tim3^+^ terminally dysfunctional cells, where it promoted their survival in tumor tissue. CXCR6 deletion resulted in decreased tumor growth control, highlighting the value of the dysfunctional state in mediating antitumor immunity and the importance of the TCF1:CXCR6-CXCL16 regulatory circuit in CD8^+^ TILs.

Our data-driven approach provides a new methodological framework for single-cell genomics analysis in many settings, including healthy tissues, cancer, and other diseases. Our approach is based on the assumption that when observing cells under different conditions, for example residing in different TMEs, each cell should manifest both shared and context-specific expression programs. With a flexible incorporation of covariates to matrix factorization, GMDF provides a robust framework to capture this structure and uncover latent expression programs (or meta-features) accordingly, allowing each cell to have more than one “context” (*e.g.*, patient sex, age, treatment status). GMDF also requires factorization reproducibility and provides a framework to identify generalizable patterns across cohorts, eliminating patterns that arise in an idiosyncratic manner or due to technical artifacts. While here we focused on expression programs shared across CD8^+^ TILs in various TMEs, context-specific expression programs could help identify ways to unleash immune responses in specific tissues, under specific conditions, or in specific cell states.

Expression of the decoupled pan-cancer dysfunction program predicted failure to respond to ICB, yet CXCR6 expression increased upon ICB in both melanoma patients (17) and in murine models. Our data showing that CXCR6 supports the maintenance of dysfunctional CD8^+^ TILs and tumor control, together with a recent report showing that CXCR6 mediates interactions with perivascular DCs to promote survival of effector cells (45), are consistent with a model where CXCR6 increase upon ICB supports the anti-tumor immune response. Our data showing that CXCR6 deletion specifically affected dysfunctional CD8^+^ TILs and compromised tumor control, positions CXCR6 as a positive regulator of the dysfunctional T cell state and supports the pliability and therapeutic potential of the dysfunctional T cell state. Our pan-cancer dysfunction program may contain additional positive regulators that support the dysfunctional T cell state, as well as negative regulators that temper the functional properties of dysfunctional T cells.

CD8^+^ T cells are known to undergo a modified differentiation trajectory in the TME. *Tcf7*/TCF1 plays a role early in this trajectory, where it restrains effector differentiation and preserves a pool of stem-like CD8^+^ T cells that seed the effector response upon ICB and are required for sustained ICB response (23,24,46). Our data show that TCF1 also represses CXCR6 expression. Thus, as stem-like CD8^+^ T cells differentiate into effector cells and lose TCF1 expression, they start to express CXCR6. CXCR6 expression further increases as cells progress along the effector differentiation trajectory and eventually become dysfunctional of “exhausted”. Our data support the hypothesis that CXCR6 acts at this later stage to preserve dysfunctional cells that can still exert a meaningful level of tumor control. Further, TCF1^+^ stem-like CD8^+^ T cells have been shown to reside in specific tumoral niches (47). This spatial positioning could be due in part to the repression of CXCR6, which has been shown to direct intra-tumoral positioning of cells (Di Pilato et al., 2021). Thus, the TCF1: CXCR6-CXCL16 axis that we uncover may orchestrate the differentiation trajectory of intra-tumoral CD8^+^ T cells by directing their spatial localization.

Unlike other chemokine ligands, CXCL16 can also be expressed as a transmembrane protein (48–50). CXCR6 binding to transmembrane CXCL16 can mediate cell adhesion (51) and reverse signal into CXCL16-expressing cells to modulate their phenotype (52). Thus, CXCR6-CXCL16 interactions could modify CXCL16^+^ APCs to promote their production of factors, such as IL-15 (45) and CX3CL1, the ligand for CX3CR1, to support the survival and maintenance of dysfunctional CXCR6^+^CD8^+^ TILs. Our data indicate that most APC populations express CXCL16 with some variability across cell types depending on the cancer being examined. However, in ovarian cancer, CXCL16 is expressed on subpopulations of malignant cells. Previous studies have shown that CXCL16 expression in ovarian cancer cells promotes their invasion and migration capabilities (53) and that CXCL16 from breast or colon cancer cells can influence T cell infiltration into the tumor (54, 55). Future studies should examine how CXCL16^+^ APC may be modified by contact with CXCR6^+^ dysfunctional CD8^+^ T cells and how CXCL16 expression might impact not only T cell recruitment, but also T cell longevity and direct cancer-T-cell interactions in the tumor tissue.

In conclusion, our data show that CXCR6: (**1**) is a robust marker of dysfunctional CD8^+^ T cells across an array of human tumor types, (**2**) is directly regulated by TCF1, and (**3**) is critical for sustaining the dysfunctional CD8^+^ T cell pool and tumor control. Our study thus reveals new aspects of T cell dysfunction and defines a regulatory axis essential for T-cell-mediated tumor control, opening new avenues for the development of biomarkers and interventions in cancer and other diseases of deficient or dysregulated immunity.

## Acknowledgments

We thank Rajesh Krishnan for cell sorting, Leslie Gaffney for help with Fig. preparation, and Geoffrey Fell, MS for statistical help. We thank the Human Tumor Atlas Network (HTAN) and Human Tumor Atlas Pilot Project (HTAPP) consortium for the ongoing support, fostering collaborations and new ideas across laboratories, teams, and institutes. This work was supported by grants from the National Institutes of Health (R01CA187975 to ACA), and by the Klarman Cell Observatory at the Broad Institute and HHMI. A.C.A. is a recipient of the Brigham and Women’s President’s Scholar Award. A.R. was an Investigator of the Howard Hughes Medical Institute. G.E. is a Cancer Research Institute Irvington Fellow supported by the Cancer Research Institute (CRI). L.J.A. is a Chan Zuckerberg Biohub Investigator and holds a Career Award at the Scientific Interface from BWF. L.J.A. was a fellow of the Eric and Wendy Schmidt postdoctoral program and a CRI Irvington Fellow supported by the CRI.

## Authors Contributions

Conceptualization: L.J., K.T., A.C.A, A.R.

Methodology: L.J., K.T., G.E., A.C.A., and A.R.

Investigation: L.J., K.T., G.E., G.D.

Formal analysis: L.J., K.T., G.E., C.L., E.G., A.M.

Resources: L.J., K.T., G.E., G.D.

Writing: L.J., K.T., A.C.A., and A.R.

Supervision: O.R.R., A.C.A., and A. R.

Funding acquisition: A.C.A. and A.R.

## Methods

### Generalized Matrix Decomposition Framework (GMDF)

Given a set of *d* datasets collected under different conditions, GMDF decomposes each of the original datasets *E_i_* (*n_i_ x m*) using latent shared (*W*) and context specific (*A_j_*) metagenes:

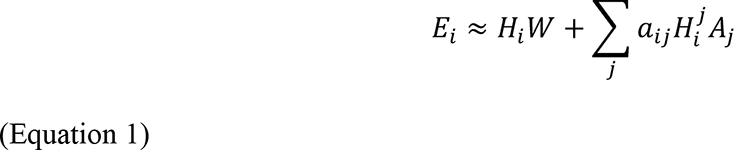

where all the factor matrices are constrained to be non-negative. 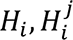 are the usage matrices of the latent programs in dataset *i*. Note that *W* (k x m) is shared across all datasets (i = 1…d), while *A_j_* (k_j_ x m) is used only if the binary parameter *a_ij_* is 1, denoting that condition *j* holds in sample *i*. A condition is defined by the user and depends on the specific structure and features of the data. It can denote the disease subtype, treatment status, data source, etc.

The optimization problem is then defined as

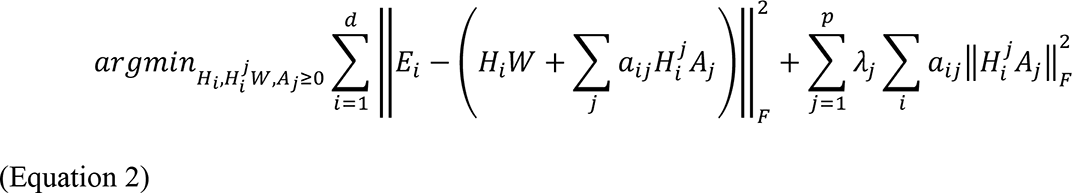

where λ_*j*_ is a tuning parameter which allows adjusting the size of context-specific effects.

### Solving GMDF with block coordinate descent

To find a solution to the non-convex GMDF optimization problems, the variables are divided into blocks. Using block coordinate descent, the objective is iteratively minimized with respect to each block, while holding the others fixed:

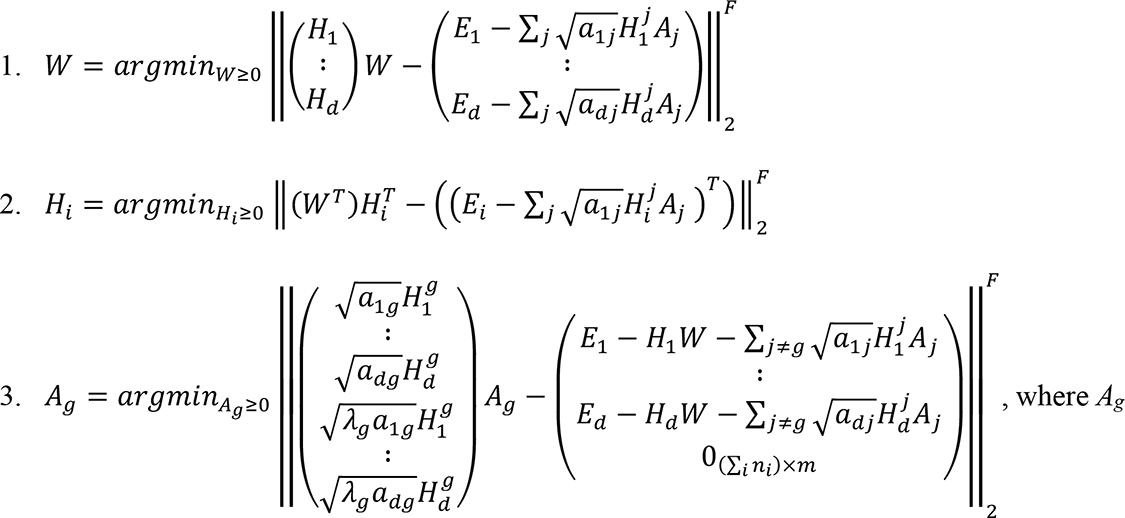

represents the programs specific to context *g*.

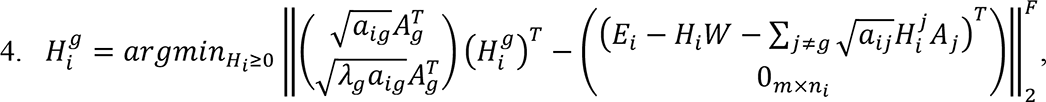

where 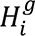 represents the usage of programs specific to context *g* in cohort *i*.

Each of the optimization subproblems above requires solving a nonnegative least-squares problem. Each of these subproblems is solved by using the fast block principal pivoting algorithm (56), implemented using the Rcpp and LIGER packages in R. Convergence criteria: After each iteration (1-4 described above), GMDF updates a convergence variable denoting the improvement in the score of the solution (computed as shown in Equation 2). The convergence variable drops semi-monotonically. Once it is below a pre-defined convergence threshold (with a default of 1*10^-4^), GMDF outputs the single run solution.

### Obtaining a consensus GMDF solution

GMDF is non-convex, and therefore has multiple non-unique optimal solutions, in addition to sub-optimal solutions that can be obtained if reaching a local minimum. To alleviate these problems and generate a single robust solution, GMDF uses random subsampling of the data, and multiple initialization conditions to generate multiple solutions. The different solutions are then aggregated such that similar solutions that were repeatedly identified across various data sampling and initializations are combined. More specifically, all the shared programs (W matrixes), are clustered via hierarchical clustering, with Euclidean distance as the metric to evaluate similarity between programs. Clusters with less than 3 metagenes are removed and the others are converted to final metagenes by computing the mean loading of each gene in each cluster. The same procedure is performed for the *A_j_* matrixes. This scheme mitigates outlier effects, reduces overfitting, and makes the final solution less dependent on the *k* and *k_j_* parameters, identifying the number of the embeddings’ dimensions directly from the data.

### Comparing GMDF to LIGER and iNMF

To compare GMDF to LIGER (26) and iNMF (25),(25)(Yang and Michailidis, 2016) data was simulated with shared and dataset-specific gene modules, where a shared gene module consists of genes that are correlated across the cells in all the datasets, and a dataset-specific module consists of genes that are correlated across the cells only in a specific dataset, where in the other datasets there is no correlation. Each dataset consisted of 200 cells, and module size was set to 100 genes. The results obtained with GMDF and LIGER/iNMF are shown in Fig. S1B-E, with the ground truth denoted by the vertical color bar to the left. As depicted in Fig. S1B-E, in contrast to LIGER/iNMF, GMDF identified all the programs and assigned them correctly as shared or specific to dataset 1 or 2. To further test LIGER, we also used another analysis mode in the LIGER package, where, beyond the decomposition, LIGER also identifies differentially expressed genes. However, also when applied to identify differentially expressed genes across the different datasets, LIGER did not identify the context-specific modules.

### Data sets and pre-processing

Processed data in the form of raw counts or transcripts per million (TPM) were obtained from the Gene Expression Omnibus (GEO), accession numbers: GSE163108 (glioblastoma (15)), GSE115978 (melanoma cohort 1 (11)), GSE120575 (melanoma cohort 2 (17)), GSE131309 (sarcoma (12)), GSE131907 (lung, Kim (13)), GSE114724 (breast cancer (10)), GSE98638 (hepatocellular carcinoma (19)), and GSE108989 (colorectal cancer (18)). Pancreatic ductal adenocarcinoma (PDAC) TIL scRNA-Seq data was obtained from the Genome Sequence Archive (GSA), accession number CRA001160, project PRJCA001063. An additional lung cancer TIL scRNA-seq data (14) was downloaded from SCope (https://gbiomed.kuleuven.be/scRNAseq-NSCLC). All data were converted to log_2_(TPM/10+1), as previously defined (22). Throughout, Overall Expression was computed as previously defined (11). The UMAPs shown in Fig. 2 and S3 were obtained with the Seurat package, using the first 30 PCs obtained with top 2,000 most variable genes identified with the *FindVariableFeatures* function (default parameters).

### GMDF application to the pan-cancer CD8 T cell cohort

GMDF was applied 25 times with different (random) initialization parameters, and random sub-sampling procedure. For each run, 200 cells were subsampled from each patient in each cohort, and the topmost variable genes were selected, with variation threshold of 0.2 in the LIGER function selectGenes. GMDF parameters were 5 for *k* and *k*’, convergence threshold 1*10^-4^, and *a_ij_* = 1 if and only if *i* = *j*, to identify shared and cohort-specific programs. Following the 25 runs, a consensus solution was obtained as described above (“**Obtaining a consensus GMDF solution”** section). Lastly, for each consensus shared program, the top 100 genes with the highest loading were used to define the pertaining program.

### Decoupling convolved expression programs

T cell “activation” and “chronic activation” scores were computed separately for each dataset. Activation scores were defined as the Overall Expression (11) of the cytotoxic markers (*NKG7, CCL4, CST7, PRF1, GZMA, GZMB, IFNG, CCL3*) minus the Overall Expression of memory/naïve markers (*CCR7*, *TCF7*, *LEF1*, *SELL*). Chronic activation scores were defined as the Overall Expression of the pan-cancer chronic activation program identified by GMDF (**Table S1**). Chronic activation scores were then plotted as a function of the activation scores using a locally-weighted polynomial regression (LOWESS, black line in Fig. 1C). The deviation from this regression line was used to classify CD8 T cells into four groups: Cells with a low activation score (below the 25^th^ percentile) were classified as naïve/memory-like cells, while the others were considered effector or dysfunctional, if their activation scores were (0.25 standard deviations) higher or lower than expected given their chronic activation scores, respectively. Cells with a high activation score (>25^th^ percentile) that fit the regression line were annotated as “balanced” and were not used in the analysis below. In each dataset, genes that were significantly overexpressed in one subset of cells (*e.g.*, dysfunctional) compared separately to each of the other two subsets (*e.g.*, dysfunction *vs*. naïve/memory and dysfunction *vs*. effector), were identified as markers of that cell subset in that dataset. These pairwise comparisons were performed with a multilevel regression model: 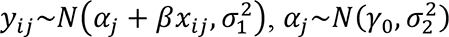, where *y_ij_* denotes the expression of the gene in cell *i* from sample *j*, *x_ij_* denotes the total log-transformed number of reads detected in cell *i* in sample *j*, and *a_j_* is the sample-specific intercept. The lme4 (57) (https://CRAN.R-project.org/package=lme4) and lmerTest R packages (58) were used to fit the model, compute p-values, and identify the latent variables that maximize the posterior probability

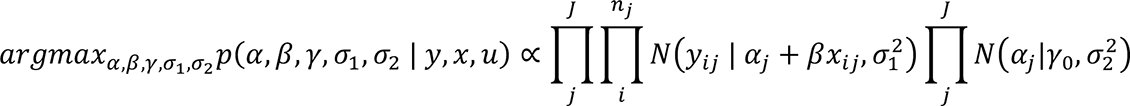

To aggregate the results across all datasets, pan-cancer p-values were obtained for each of the three pairwise comparisons using Fisher’s combined probability test. Genes that were repeatedly identified as markers of one of the three states across all datasets and had a combined p-value < 1*10^-10^ were denoted as pan-cancer markers of the pertaining state.

### CXCL16 pan-cancer macrophage program

scRNA-Seq datasets of macrophages from melanoma (11, 17), sarcoma (12), and glioblastoma (15) were analyzed to identify genes co-expressed with CXCL16. In each dataset genes co-expressed with CXCL16 were identified using partial Spearman correlation, accounting for the total number of genes detected in each cell. The CXCL16 program (**Table S4**) includes genes significantly co-expressed with CXCL16 (t-test, Benjamini-Hochberg FDR < 0.05) in all four datasets (Fisher’s combined probability, Benjamin-Hochberg FDR < 1*10^-3^). In addition, in each dataset the gene ranks based on the partial Spearman correlation analyses were used as input for gene set enrichment analysis (GSEA). The latter was preformed using the fast GSEA package to obtain the normalized enrichment scores (NES) and plots shown in Fig. 3B, using the KEGG_ANTIGEN_PROCESSING_AND_PRESENTATION gene set.

### Mice

6–8-week-old male or female C57BL/6 (Stock No. 000664), CXCR6 KO (Stock No. 005693), OT-1 (Stock No. 003831), Cas9 (Stock No. 028555), and CD45.1 (Stock No. 002014) transgenic mice were purchased from the Jackson Laboratory. CXCR6 KO mice were crossed to OT-1. All mice were housed in a vivarium under SPF conditions, in cages of up to five mice, and fed a special rodent diet. All experiments involving laboratory animals were performed under protocols approved by the Harvard Medical Area Standing Committee on Animals (Boston, MA) and followed IACUC guidelines on the ethical care and use of animals.

### Mouse tumor cell lines

B16F10 was purchased from ATCC. MC38Ova^Hi^ was generously provided by Dr. Nick Haining and B16Ova was kindly provided by Dr. Kai Wucherpfennig. B16F10 and B16Ova were grown in RPMI-10% FBS at 37°C and injected subcutaneously into the flanks of mice at 3-4×10^5^ cells per mouse. MC38Ova^Hi^ cells were grown in DMEM-10% FBS at 37C and injected subcutaneously into the right flank of mice at 1×10^6^ cells per mouse.

### Tumor experiments with immune checkpoint blockade

Tumor size was measured in two dimensions by caliper and is expressed as the product of two perpendicular diameters. In immune checkpoint blockade treatment experiments, B16Ova tumors were treated with a combination of anti-PD-L1 (10F.9G2) (200μg per mouse) and anti-Tim-3 (RMT3-23) (200μg per mouse) antibodies or control immunoglobulin (Rat IgG2a and Rat IgG2b) i.p. on days specified for each experiment. Mc38Ova^Hi^ tumors were treated with anti-PD1 (RMP1-14) (100μg per mouse) i.p. on days specified for each experiment. Mice were then monitored every two days for tumor growth.

### TIL isolation

TILs were isolated by dissecting the tumor mass and mincing the tumor tissue prior to digestion with Collagenase D (2.5 mg/ml) for 20 minutes at 37°C. Tumors were then dissociated to single cell suspensions by passing tissue through a 70μM filter prior to centrifugation and analysis by flow cytometry. For cytokine analysis, tumors were dissociated as above and cells underwent a density separation via percoll gradient before being plating for *ex vivo* stimulation.

### Flow cytometry

Single cell suspensions were stained with antibodies against surface molecules. Antibodies against CD8 (53-6.7), CD45.1 (A20), CD45.2 (104), CCR7 (4B12), CD103 (2E7), PD-1 (RMP1-30), PD-1 (29F.1A12), Ly-6C (HK1.4), CD11c (N418), I-A/I-E (M5/114.15.2), CD39 (Duha59), CD64 (X54-5/7.1), CD90.1 (OX7), CX3CR1 (SA011F11), CD107a (1D4B), Lag3 (C9B7W), TCRβ (H57-597), Tim3 (5D12), XCR1 (ZET), and CXCR6 (SA051D1) were purchased from BioLegend. The antibody against Tigit (GIGD7) was purchased from eBioscience. Antibodies against CD11b (M1/70), CD24 (M1/69), CD45 (30-F11), CD8a (53-6.7), human-NGFR (C40-1457), TCRb (H57-597), Thy1.1 (OX7), and Tim3 (5D12) were purchased from BD Biosciences. The antibody against F4/80 (CI:A3-1) was purchased from BioRad. Annexin V (BD) staining was performed per manufacturer’s protocol. Fixable viability dye Zombie UV (Biolegend) or 7AAD (Biolegend) were used to exclude dead cells.

For intracellular staining, eBioscience Foxp3/transcription factor staining buffer set was used per manufacturer’s protocol. The antibody against Bcl-2 (BCL/10C4) was purchased from Biolegend. The antibodies against TCF1 (C63D9) and Bim (C34CS) were purchased from Cell Signaling. The antibody against Tox (REA473) was purchased from Miltenyi Biotec, and Ki-67 (SolA15) was purchased from eBioscience.

For intra-cytoplasmic cytokine (ICC) staining of TILs, cells were stimulated *ex vivo* with 5 μg/mL OVA257-264 peptide (GenScript) for 4 hours in the presence of Golgi stop (BD Biosciences) and Golgi Plug (BD Biosciences) prior to cell surface and ICC staining. Importantly, the antibody detecting CD107a (1D4B) was added to the cells during stimulation. Following fixation and permeabilization, staining with antibodies against the following was performed: IL-2 (JES6-5H4), TNF-a (MP6-XT22), IFN-g (XMG-1.2), and Granzyme B (GB11) which were purchased from Biolegend.

For intracellular staining of CXCL16 (12–81) from BD, cells from tumors were cultured *ex-vivo* from tumors for 4 hours in the presence of Golgi Plug (BD Biosciences) prior to cell surface and intracellular staining.

All data were collected on a BD Symphony (BD Biosciences) or Fortessa (BD Biosciences) and analyzed with FlowJo 10.7.1 software (TreeStar).

### Adoptive cell transfers

Tumor cells were injected into the flanks of recipient mice as outlined above. On day 6 post tumor implantation, mice were randomized by tumor size and given T cells suspended in PBS via tail intravenous injections. For CRISPR-KO experiments, 3×10^6^ (for B16Ova) or 2.25×10^6^ (for Mc38Ova^Hi^) CRISPR CXCR6-KO or CRISPR-Control transduced cells were transferred. For overexpression experiments, 5×10^5^ CXCR6-OE or Control transduced cells were transferred. For CXCR6 KO experiments, 7.5×10^5^ CXCR6-KO or Control cells were transferred. Mice were then followed every two days and tumor sizes were measured until the day of tumor harvest for subsequent isolation of TILs and analysis via flow cytometry.

### Purification, lentiviral transduction, and *in vitro* culture of primary CD8+ T cells for adoptive cell transfer experiments

CD8^+^ T cells from spleens and lymph nodes of CD45.2 Cas9+ OTI+ mice (for CRISPR/Cas9 KO transfers) or CD45.2 OTI+ mice (for OE transfers) were isolated using CD8a microbeads (Miltenyi) and plated at a concentration of 5×10^5^ cells per well in the presence of IL-2 (6ng/mL) in 24-well plates previously coated with 1μg/mL anti-CD3/anti-CD28. The following day, cells were transduced at an MOI of 100 with lentiviral vectors. For CRISPR/Cas9 KO experiments, the vector contained an sgRNA targeting CXCR6 and a Thy1.1 gene to be used as a marker for transduction. A vector lacking the sgRNA but containing Thy1.1 was used as a control. For OE experiments, cells were transduced with a bidirectional lentiviral vector encoding CXCR6 and hNGFR (CXCR6 OE) or Thy1.1 or hNGFR (Control). Cells were propagated in culture in the presence of IL-2 for 6 days. On the day of transfer, cells were harvested from the plates and stained with an antibody against Thy1.1 (for KO experiments) or hNGFR (for OE experiments) and FACs sorted to purify highly transduced cells.

For CXCR6 KO adoptive transfer experiments, cells were isolated from CD45.2 CXCR6 KO OTI+ mice or control CD45.2 OTI+ mice and plated/activated as above. Cells were activated for 48 hours before replating and allowing to propagate in culture in the presence of IL-2 for a total of 6 days. On the day of transfer, cells were harvested from the plates and transferred into mice.

### sgRNA design and testing

sgRNA guides were designed using the Broad Institute’s GPP sgRNA designer portal (https://portals.broadinstitute.org/gpp/public/analysis-tools/sgrna-design). Three candidate sgRNAs were chosen and each cloned into an sgRNA plasmid using BsmBI sites (Addgene #85453, modified with Thy1.1 in place of Tdtomato). Candidate sgRNA plasmids were transfected into doxycycline-inducible-Cas9 expressing murine 3T3 cells. After 48 hours, transfection efficiency was determined and gDNA was collected. gDNA was used to perform a TIDE (Tracking of Indels by DEcomposition) assay (59). Briefly, PCR was used to amplify a roughly 700bp region around the expected sgRNA binding site. The PCR product was purified by agarose gel purification and Sanger sequenced. The resulting chromatogram was uploaded to the TIDE website for analysis and each sgRNA was evaluated compared to a control-transfected sample to determine the guide editing efficiency. The guide with the highest editing efficiency was chosen to move forward for lentivirus production (CXCR6 guide: 5’-GCAGAGTACAGACAAACACC).

### Plasmid construction and lentivirus production

The Thy1.1 tagged, sgRNA containing plasmid was described previously in the **Methods** (**sgRNA design and testing**). The bidirectional CXCR6 overexpression vector was generated by cloning the CXCR6 cDNA amplified from Genscript Accession No. NM_030712.4 by PCR into the BdLV as previously described (60) using AgeI and SalI. PCR primers for amplification were as follows: Fw-primer: 5’-AAAACCGGTGCGCCACCATGGATGATG; Rev-primer: 5’-AAAGTCGACACACTGGACTAGTGGATCCCT.

VSV-pseudotyped third-generation lentiviruses were made as described previously (61). Briefly, 293T cells were transiently co-transfected with five plasmids, then underwent a media change 12 hours post-transfection. Cell supernatant was collected 26-30 hours later and virus was purified by ultracentrifugation. Viral titers were determined on 293T cells by limiting dilution.

### *In vitro* time-course of T cell activation and bulk RNA-Seq

CD8^+^ T cells from spleens and lymph nodes of wildtype OTI+ mice and E8i-Cre+ TCF7^FL/FL^ mice (n=3 per group) were isolated using CD8 microbeads (Miltenyi). After taking a timepoint 0 sample, cells were plated on plates previously coated with 1 μg/mL anti-CD3/anti-CD28 in the presence of murine IL2 (6ng/mL) and cultured at 37°C. For each timepoint, 1,000 cells were sorted into 5μL of Buffer TCL (QIAGEN) supplemented with 1% 2-mercaptoethanol. Samples were processed using the SMART-Seq2 protocol (62), and sequenced on an Illumina NextSeq.

RNA-seq reads were aligned using STAR to mouse genome version mm10, and expression levels were calculated using RSEM (Li and Dewey, 2011) using annotated transcripts (mm10), followed by further processing using the Bioconductor package DESeq2 in R(Anders and Huber, 2010). Data was normalized using TPM normalization, and differentially expressed genes were defined using the differential expression pipeline on the raw counts with a single call to the function DESeq2 (reported p-values obtained via Wald test only for Benjamin-Hochberg FDR< 0.05).

### ChIP-PCR

CD8^+^ T cells from spleens and lymph nodes of TCR-transgenic E8i-Cre+ TCF7^FL/FL^ mice and TCR-transgenic E8i-Cre-TCF7^FL/FL^ mice were isolated using CD8 microbeads (Miltenyi). Cells were fixed for 10 minutes at room temperature with 1% formaldehyde and quenched with glycine. Cells were lysed with an SDS-lysis buffer prior to chromatin shearing via sonication. Sheared chromatin was “pre-cleared” using Protein A Agarose/Salmon Sperm DNA beads from Sigma-Aldrich. Pre-cleared chromatin from the equivalent of 3.57×10^6^ cells was used in the chromatin IP reactions along 3μg of either anti-TCF1 (C63D9) or Normal Rabbit IgG, both purchased from Cell Signaling. Protein-chromatin complexes were then bound with Protein A Agarose/Salmon Sperm DNA beads and underwent multiple rounds of washes. Finally, complexes were eluted off the beads, underwent reverse crosslinking with NaCl, and treated with proteinase K before isolating DNA using the QIAquick PCR Purification kit from Qiagen. DNA content from chromatin IPs was assessed using SYBR Green amplification with the following primers designed from the promoter region of the murine CXCR6 gene: Fw-primer: 5’ – GAGGCAGACCTTTAGTGAGCA– 3’. Rev-primer: 5’ – TAGCTCGCACCGTATACACA – 3’. Results from Chromatin IP reactions were normalized to their “input” samples, or sample taken prior to Chromatin IP.

### Mouse single cell RNA-seq

E8i-Cre-TCF7^FL/FL^ (wildtype) and E8i-Cre^+^ TCF7^FL/FL^ (*Tcf7* cKO) mice were injected with 2.5×10^5^ B16Ova cells subcutaneously in their flanks. On day 13, tumors were harvested (n=3 per group) and dissociated to single cell suspensions prior to FACS. CD45^+^ TCRb^+^ CD8^+^ T cells were sorted (2,000 cells/mouse). Cells from mice from the same group were pooled and loaded for encapsulation for on the Chromium system (10x Genomics). Libraries were prepared using 5’ sequencing 10X Genomics kit v1 according to the Manufacturer’s protocol. Libraries were sequenced on an Illumina HiSeq.

### Mouse single cell RNA-seq analysis

Gene counts were obtained by aligning reads to the mm10 genome using CellRanger software (v1.3 10 3 Genomics). Cells were removed if they contained >5% mitochondrial transcripts or <200 genes, retaining 309 cells with 14,215 detected genes for further analysis. A log-transformed normalized count matrix was used as input for Principal Component Analysis (PCA) and the top 16 PCs were kept based on a drop in the proportion of variance explained by subsequent PCs. We confirmed that the resulting analyses were not particularly sensitive to this choice. Cells were clustered based on their top 16 PCs scores with the Louvain-Jaccard graph clustering algorithm (63), as previously described (64, 65) using the FindNeighbors (*k*=20) followed by FindClusters (resolution=0.5) functions with default parameters. Analysis and plots were generated using Seurat package (version 4.0.1) for R with default parameters.

## QUANTIFICATION AND STATISTICAL ANALYSIS

Significant differences between two groups were analyzed using GraphPad Prism 8 using paired or unpaired two-tailed Student’s t test. Tumor growth curves were analyzed using linear mixed effects models to test the trajectory of growth between various genotypes or treatments over time controlling for mouse. In mouse RNA-Seq data, differentially expressed genes following RNA-seq were defined using the differential expression pipeline on the raw counts with a single call to the function DESeq (p-values obtained via Wald’s test and corrected for multiple hypotheses testing using Benjamin-Hochberg correction with a 0.05 FDR cutoff). scRNA-Seq data was analyzed as describe above, with mixed-effect models used to account for confounders when examining differential expression of genes and gene programs. Values of *p < 0.05, **p < 0.01, ***p < 0.001 and **** p<0.0001 were considered statistically significant.

## Supplementary Figure Legends

**Figure S1.**
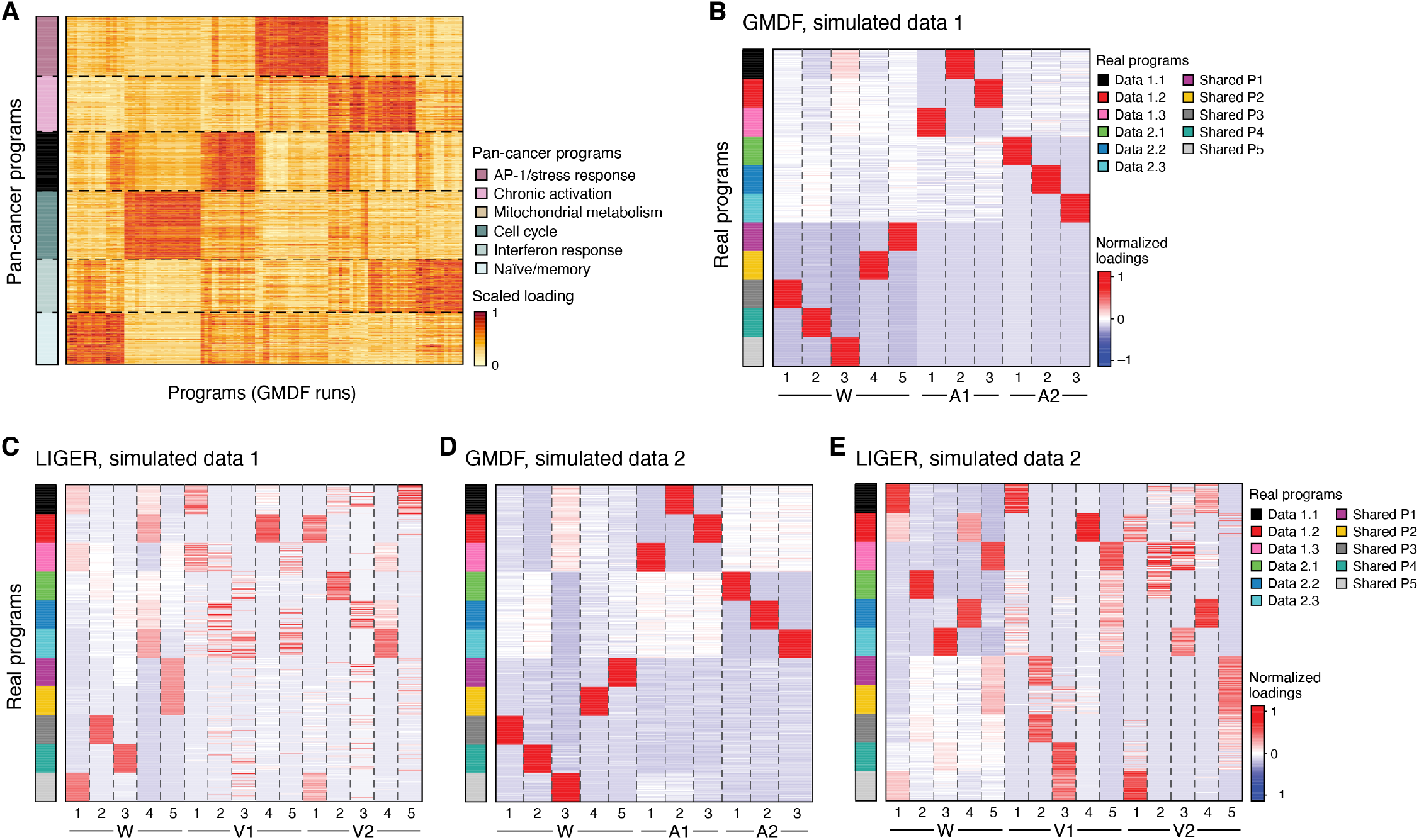
Comparison of GMDF to LIGER on simulated data. **(A)** GMDF identified 6 pan-cancer expression programs in T cells. Scaled loadings (colorbar) for top 100 genes (rows) from each program (right colorbar) across GMDF solutions (columns). **(B-E)** GMDF outperformed LIGER in recovering shared and context specific programs on simulated data. Normalized loadings (color bar) of the dataset genes (rows) across the final shared (W) and context-specific (A1 and A2, or V1 and V2) metagenes (columns) identified by GMDF (B,D) and LIGER (C,E) on two datasets (1 (B,C) and 2 (D,E)) simulated with predefined shared and dataset-specific metagenes (left “real programs” colorbar).

**Figure S2.**
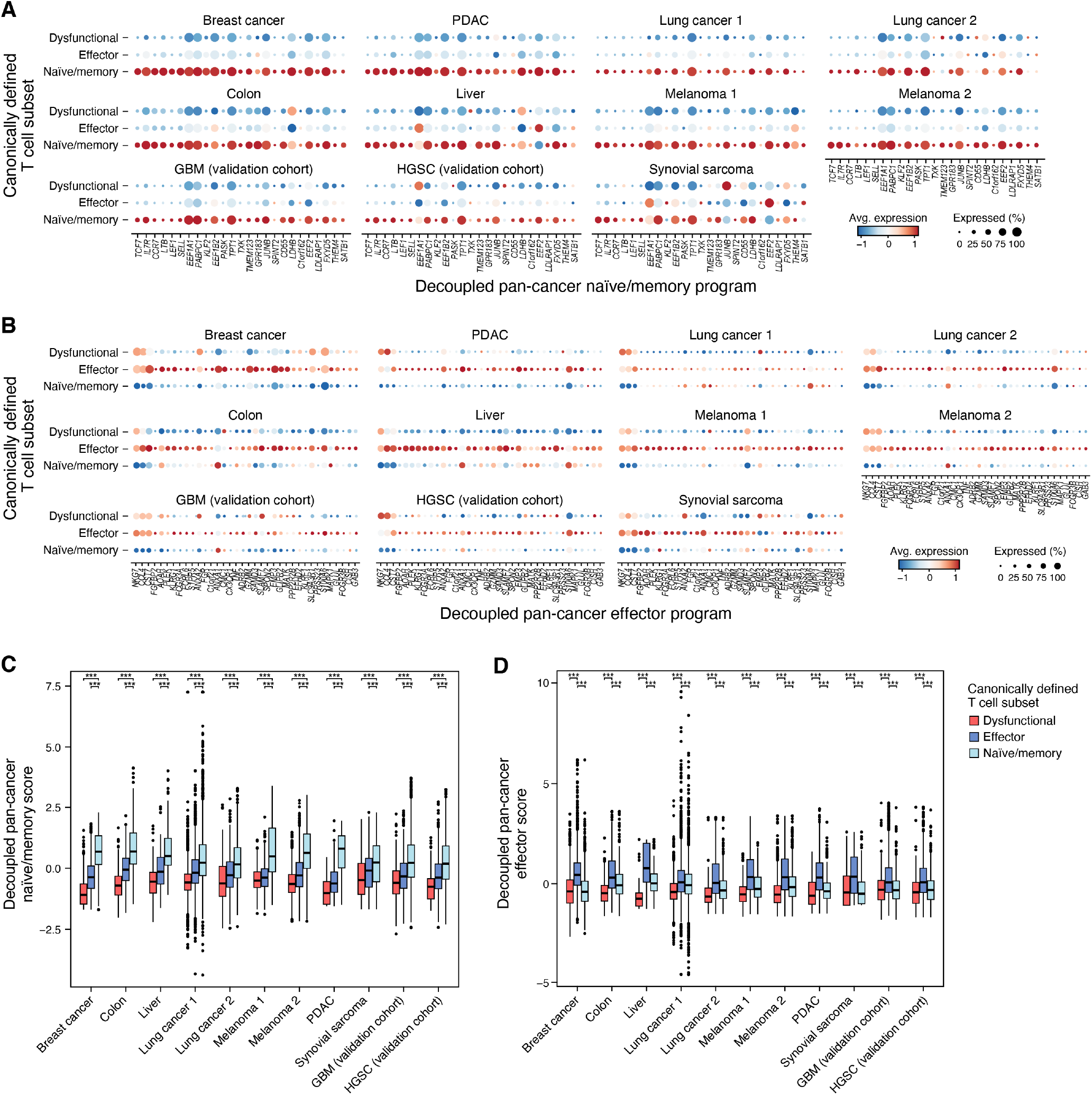
Generalizable pan-cancer features of chronic T cell activation. **(A, B)** CD8^+^ T cells from 11 tumor scRNA-seq studies (x axis) were stratified by expression of canonical markers (naïve/memory (light blue): *CCR7, TCF7, LEF1, SELL;* effector (blue): *NKG7, CCL4, CST7, PRF1, GZMA, GZMB, IFNG, CCL3;* dysfunctional (red): *PDCD1, TIGIT, HAVCR2, LAG3, CTLA4*). These canonical markers were then removed from the decoupled pan-cancer naïve/memory program (A) or the decoupled pan-cancer effector program (B) (to avoid circularity), and their Overall Expression scores were computed (**STAR Methods)**. Distribution of these Overall Expression scores (y axis) of the decoupled pan-cancer naïve/memory (A) and effector (B) programs are shown across the different cell subsets. Middle line: median; box edges: 25^th^ and 75^th^ percentiles, whiskers: most extreme points that do not exceed ±IQR*1.5; further outliers are marked individually. **(C, D)** Mean expression (dot color, color bar) and proportion of expressing cells (dot size) of genes (columns) in the decoupled pan-cancer naïve/memory program (C) and the decoupled pan-cancer effector program (D) in CD8^+^ T cells stratified as in (A and B). ***p < 0.001, mixed effects test.

**Figure S3.**
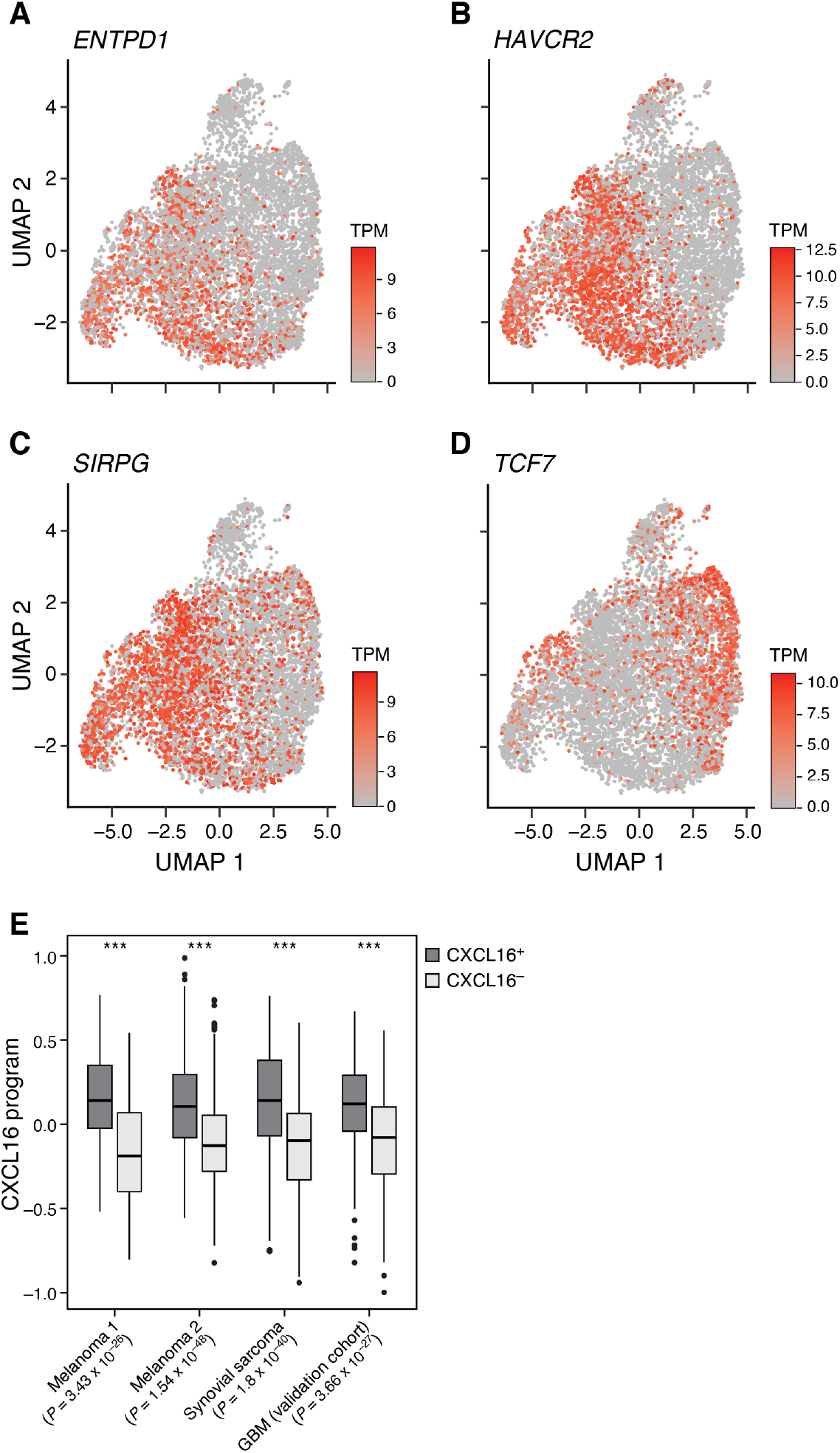
Expression of key T cell dysfunction and stemness genes in CD8^+^ TILs from patient melanomas. UMAP of CD8^+^ melanoma TILs (17) profiles colored by expression (TPM, colorbar) of T cell dysfunction genes *ENTPD1* **(A)**, *HAVCR2* **(B)**, and *SIRPG* **(C)**, or the stemness regulator *TCF7* **(D)**. **(E)** Overall Expression (y axis, **Methods**) of the macrophage CXCL16 program in CXCL16^+^ and CXCL16^-^ macrophages in the indicated human cancers.

**Figure S4.**
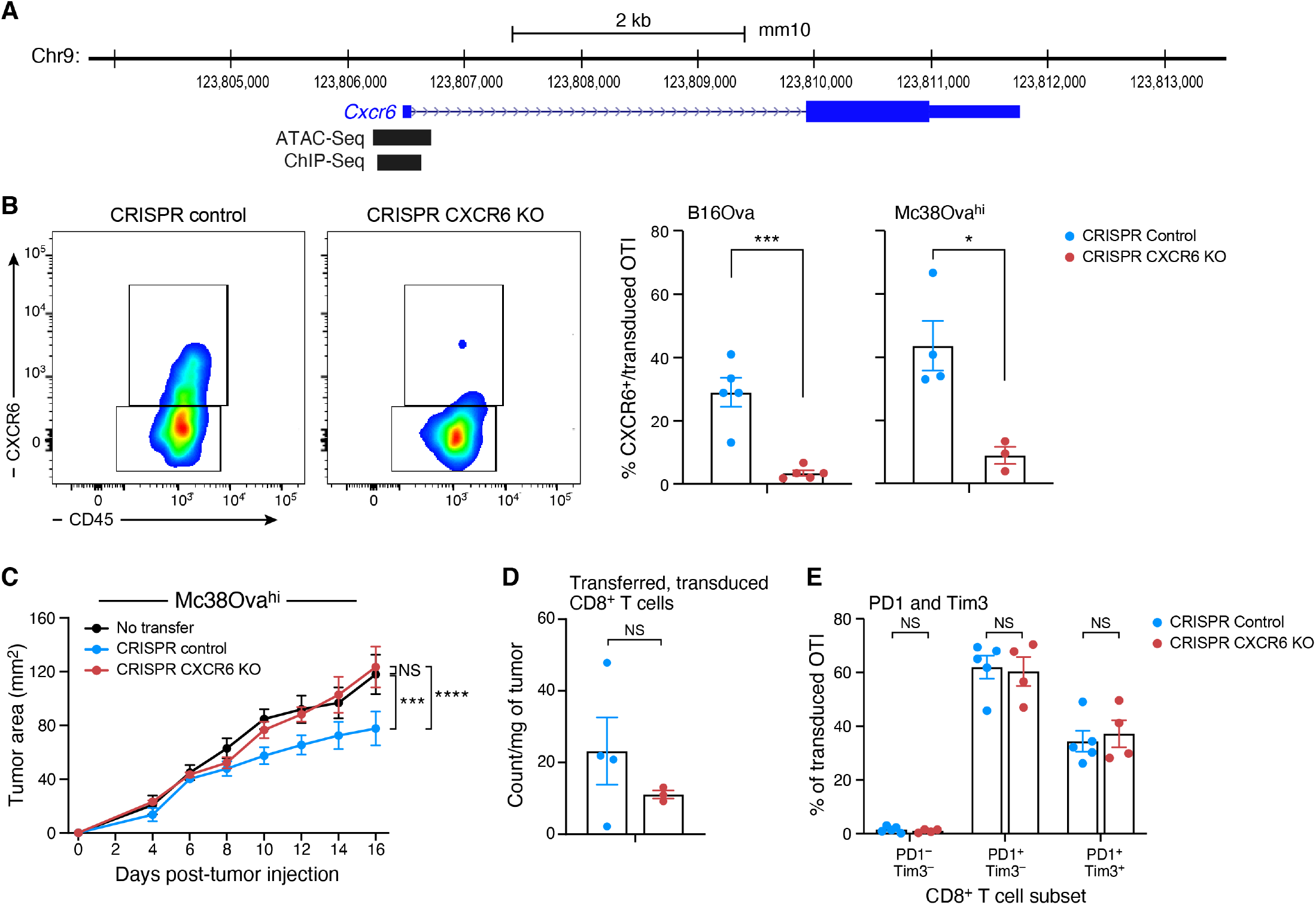
Map of CXCR6 locus and effects of CRISPR CXCR6-knockout effects on CD8^+^ T cell functionality. **(A)** Representation of the murine CXCR6 locus on chromosome 9. Blue thick boxes denote exons, thin dotted line denotes introns. Locations of ChIP-Seq peak for TCF1 in mature CD8^+^ T cells and an ATAC-Seq peak in naïve CD8^+^ T cells are shown. **(B)** Effective CXCR6 knockout by CRISPR-Cas9. Left: Representative plots of the distribution of CXCR6 (*y* axis) vs. CD45 (*x* axis) expression in control (leftmost) or CXCR6 CRISPR-KO (second from left) transferred, transduced cells from B16Ova tumors (left). Right: Percent of CXCR6 expressing cells (y axis, mean ± SEM) from control (blue) or CXCR6 CRISPR-KO (red) transferred, transduced cells from B16Ova and Mc38Ova^Hi^ tumors. **(C)** Reduced tumor control in CXCR6 CRISPR-KO in Mc38Ova^Hi^ model. Tumor size (y axis, mm^2^, mean ± SEM) over time (x axis, days) for Mc38Ova^Hi^ in Cas9^+^, CD45.1/.2 mice adoptively transferred on day 6 with CRISPR CXCR6-KO (red), CRISPR-Control (blue) OT-I T cells, or no transfer control (black). ***p < 0.001, ****p<0.0001, linear mixed model. **(D,E)** CXCR6 CRISPR-KO does not impact the numbers of transduced cells or in the PD1-and Tim3-expression profile. Number of transferred, transduced cells per mg of Mc38Ova^Hi^ tumor tissue (C, y axis, mean ± SEM) and proportions of PD1^-^ Tim3^-^, PD1^+^ Tim3^-^, and PD1^+^ Tim3^+^ transferred, transduced cells (D, y axis, Mean ± SEM) in CRISPR CXCR6-knockout (red) or CRISPR-Control (blue) OT-I T cells at time of harvest in B16Ova.

**Figure S5.**
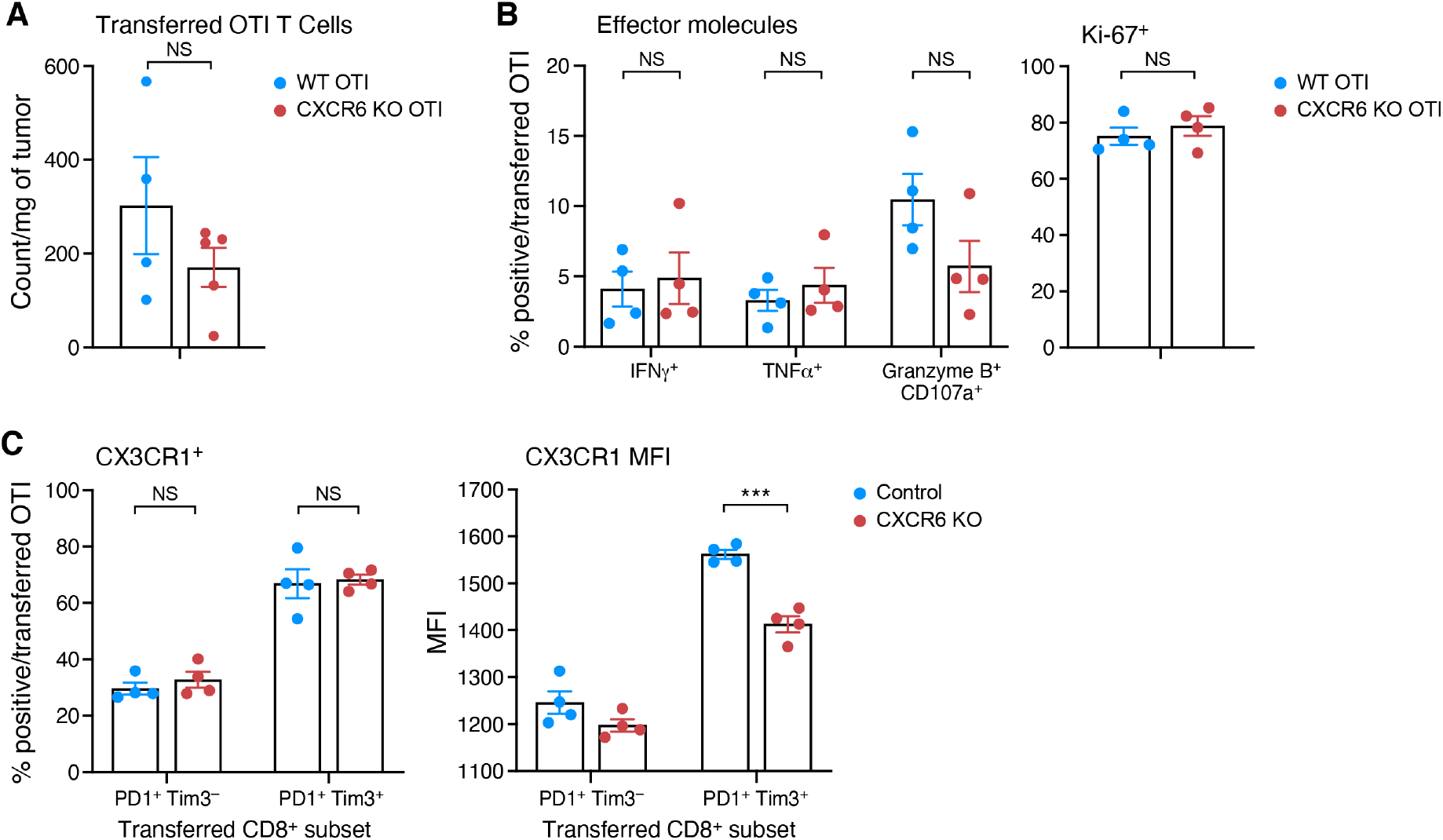
CXCR6 knockout effects on CD8^+^ T cell functionality. **(A)** CXCR6 knockout does not affect the number of transferred cells present in the tumor. Number of transferred cells per mg of tumor tissue (y axis, Mean ± SEM is shown) from control (blue) or CXCR6 KO (red) OTI cells found in B16Ova tumors. **(B)** CXCR6 KO does not affect the proliferation, cytokine production, and cytotoxicity of transferred cells. Frequency of transferred cells (y axis, mean ± SEM) that express IFN*γ*^+^, TNF*α*^+^, or Granzyme B^+^, CD107a^+^ after ex-vivo activation with Ova 257-264 peptide (left, x axis), or Ki-67^+^ (right) among control (blue) or CXCR6 KO (red) OTI cells. **(C)** Decreased expression of CX3CR1 in CXCR6 KO PD1^+^Tim3^+^ cells. Frequency of CX3CR1^+^ cells (left, y axis, Mean ± SEM) and geometric mean fluorescence (MFI) of CX3CR1^+^ cells (right y axis, Mean ± SEM) among transferred cells within PD1^+^Tim3^-^and PD1^+^Tim3^+^CD8^+^TILs. *p < 0.05, **p < 0.01, ***p < 0.001, ****p<0.0001, unpaired t test.

## Supplementary Tables

**Table S1. Six pan-cancer programs**

**Table S2. Enrichments for pan-cancer programs**

**Table S3. Uncoupled pan-cancer naïve-memory, effector, and dysfunction programs.**

**Table S4. CXCL16 program identified in macrophages.**

